# NoiTox: A Machine Learning Strategy for Peptide Toxicity Prediction Reveals Label Noise in Benchmark Datasets

**DOI:** 10.64898/2026.10.08.757741

**Authors:** Gustavo O. Almeida, Matheus C. Albuquerque, Bruno M. Silva, Olavo A. B. Silva, Ricardo S. Ferreira, Wagner Meira, Douglas V. Pires, Wallace Mateus Prata, Sabrina A. Silveira

**Author notes:** The authors wish it to be known that, in their opinion, the first two authors should be regarded as Joint First Authors.

## Abstract

**Motivation:** Peptide toxicity remains a major bottleneck in drug development, as small sequence variations can drastically alter cytotoxic profiles. Experimental assessment is costly and time-consuming, motivating computational predictors. Existing approaches often rely on single black-box architectures, limiting interpretability and obscuring the contribution of individual sequence features. Furthermore, benchmark datasets frequently define the negative class through annotation filters rather than direct experimental evidence, introducing structural label noise that can compromises training and evaluation.

**Results:** We present NoiTox, a multi-model strategy for peptide toxicity prediction integrating three complementary architectures, evaluated as four model variants: NoiToxML (Bo/full), an Extremely Randomized Trees classifier with Boruta-selected or complete physicochemical, compositional, and atomic descriptors; NoiToxDP, a deep learning model leveraging ESM-2 protein language model embeddings; and NoiToxGP, a graph neural network that combines ESM-2 residue embeddings, a fully connected residue graph with attention, and the Boruta descriptors as global features. Averaged over five random seeds, the models approached the state-of-the-art predictor ToxiPep on CD-HIT-controlled test sets, and on an independent dataset the interpretable NoiToxML variants and a probability-averaging consensus of the four models exceeded it, showing that physicochemical descriptors are highly informative for toxicity prediction. Cross-referencing negative-labeled sequences against curated databases of hemolytic, cytotoxic, and neurotoxic peptides flagged a small set annotated as toxic elsewhere, consistent with label noise by omission. However, removing the sequences that all models contradicted did not improve, and for some models reduced, ranking performance, indicating that such sequences carry useful information and are better treated as candidates for curation than discarded automatically.

**Availability and implementation:** https://github.com/goalmeida05/NoiTox.

## Introduction

The development of peptide- and protein-based therapeutics has expanded considerably in recent decades due to their high molecular selectivity, functional diversity, and potential biocompatibility when compared to traditional small-molecule drugs (Wang et al., 2022; Vrbnjak and R., 2024). Currently, more than 80 peptide-based drugs have been approved worldwide for applications in areas such as oncology, endocrinology, infectious diseases, and immunomodulation (Mengyang et al., 2025). Despite these advances, toxicity remains a major barrier to the successful clinical translation of bioactive peptides, as these molecules may induce hemolysis, cytotoxicity, immunogenic responses, or pharmacokinetic instability (Otvos and Wade, 2014; Pereira et al., 2024). Among these compounds, antimicrobial peptides (AMPs) have attracted particular attention due to their broad-spectrum antimicrobial activity; however, many AMPs exhibit limited selectivity, resulting in a narrow therapeutic window in which antimicrobial activity may be accompanied by toxicity toward host cells (Ma et al., 2024b). Structural studies have further demonstrated that small conformational changes can significantly alter peptide toxicity and selectivity, highlighting the importance of understanding structure–toxicity relationships during peptide design (Gagat et al., 2024).

Several computational approaches have been developed to predict peptide toxicity and support the design of safer therapeutic candidates. Methods such as ToxinPred2 (Rathore and others, 2024), CSM-Toxin (Morozov et al., 2023), PepNet (Han et al., 2024), and ToxiPep (Guan et al., 2025) have demonstrated promising predictive performance by employing deep learning architectures or structural modeling strategies. However, these approaches typically rely on a single architecture, and those based on deep-learning embeddings offer limited insight into which physicochemical or structural features drive toxicity predictions. Furthermore, the evaluation of these methods has been largely conducted on benchmark datasets whose negative class, sequences annotated as non-toxic, was assembled through database filters rather than direct experimental evidence, a distinction with non-trivial consequences for model training and evaluation.

To address these limitations, we propose NoiTox, a multi-model strategy for peptide toxicity prediction built around three complementary architectures, evaluated as four model variants. First, an Extremely Randomized Trees (ERT) classifier is trained on physicochemical and compositional descriptors, evaluated both on the complete descriptor set (NoiToxML full) and on a compact, biologically interpretable subset identified via Boruta feature selection (NoiToxML Bo). Second, a deep learning model (NoiToxDP) leverages ESM-2 protein language model embeddings to capture sequence-level representations learned from evolutionary context at scale. Third, a graph neural network (GNN, NoiToxGP) combines ESM-2 embeddings with a fully connected residue graph to exploit both residue-level semantics and long-range residue–residue message passing. All four models are evaluated on established benchmark datasets and compared with state-of-the-art predictors: the NoiTox variants approach the best baseline on CD-HIT-controlled test sets and, together with a probability-averaging consensus, exceed it on an independent dataset.

Beyond predictive accuracy, we present a systematic analysis of sequences consistently misclassified across all models. Cross-referencing with curated hemolytic, cytotoxic, and neurotoxic databases flagged a set of sequences labeled as non-toxic that are annotated as toxic elsewhere, which we treat as candidates for label noise by omission, a structural bias in benchmark datasets that can affect the training and evaluation of any toxicity predictor built upon them. Removing the sequences contradicted by all models did not improve performance, indicating that they carry useful information and are better handled as candidates for curation.

## Materials and methods

This section describes the NoiTox strategy for peptide toxicity prediction. We detail the data collection and curation pipeline, the encoding strategies employed across four models, an Extremely Randomized Trees classifier (Full and Boruta), an ESM-2 deep learning model, and a graph neural network, and the experimental evaluation protocol used to benchmark performance against state-of-the-art methods.

### Dataset preparation

To construct a robust dataset for model development, toxic peptides were collected from widely used benchmark datasets, including ToxinPred2 (Rathore and others, 2024) and ATSE (Wei et al., 2021), as well as experimentally validated entries from DRAMP 4.0 (Ma et al., 2024a) and dbAMP 3.0 (Yao et al., 2024). Non-toxic peptide sequences were obtained from the curated UniProt (Consortium, 2018) and Swiss-Prot (Boeckmann et al., 2003) databases. Sequences containing ambiguous or non-standard amino acid residues were removed during preprocessing to ensure descriptor consistency.

Sequence redundancy was reduced using the CD-HIT algorithm (Li and Godzik, 2006) at two sequence identity thresholds (0.9 and 0.8), enabling comparison with previous studies while also providing a more stringent evaluation scenario. The resulting dataset composition for each clustering threshold is summarized in Table 1.

**Table 1.**
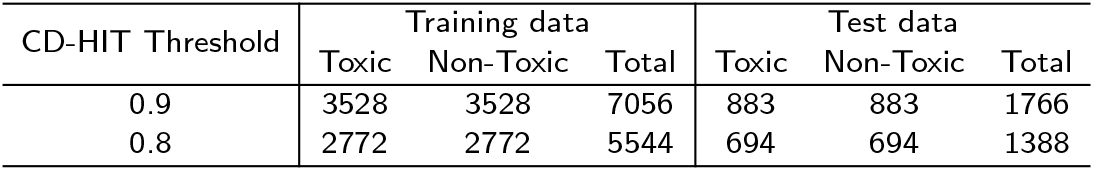
Composition of the balanced datasets generated after redundancy reduction using CD-HIT at different sequence identity thresholds.

| CD-HIT Threshold | Training data |  |  | Test data |  |  |
| --- | --- | --- | --- | --- | --- | --- |
|  | Toxic | Non-Toxic | Total | Toxic | Non-Toxic | Total |
| 0.9 | 3528 | 3528 | 7056 | 883 | 883 | 1766 |
| 0.8 | 2772 | 2772 | 5544 | 694 | 694 | 1388 |

To prevent data leakage and ensure realistic model evaluation, peptide sequences were grouped by similarity prior to splitting the data into training and testing sets using Levenshtein distance–based clustering (Berger et al., 2021; Yu et al., 2024). Additionally, an independent test set composed of experimentally validated toxic peptides from the DBAASP database (Pirtskhalava et al., 2021) was used to further assess the generalization capability of the proposed model. According to (Guan et al., 2025), who originally assembled this benchmark, the 479 positive entries are peptides with experimentally measured hemolytic activity, classified as toxic based on established hemolysis thresholds. The corresponding 479 negative entries, however, were curated from reviewed UniProt and Swiss-Prot entries by excluding sequences carrying toxin-related annotations, an absence-of-annotation criterion rather than direct experimental confirmation of non-toxicity. This asymmetry between experimentally validated positives and annotation-filtered negatives mirrors the label-noise-by-omission concern examined throughout this study, and should be taken into account when interpreting specificity values on the independent dataset (Table 3).

To further characterize the nature of toxicity predicted by the model, a validation was performed using experimentally validated datasets representing specific toxicity subtypes. Since the binary toxic/non-toxic classification adopted in model training is inherently general, this subtype-level validation aimed to assess whether the model captures biologically meaningful toxic signals beyond the broad categorical label. Four curated databases were employed for this purpose:

- DBAASP: peptides with experimentally validated cytotoxic and/or hemolytic activity were retrieved from the DBAASP database (Pirtskhalava et al., 2021), filtering for sequences between 2 and 50 amino acid residues in length and selecting entries annotated with hemolytic or cytotoxic biological activities. DBAASP is a comprehensive repository of antimicrobial peptides with detailed functional annotations, making it a valuable source for independent validation of membrane-disrupting toxic mechanisms.
- Hemolytic activity: peptides with experimentally validated hemolytic and non-hemolytic activity were retrieved from the Hemolytic2 database (Dimitrov et al., 2026), a manually curated resource containing over 13,200 fully annotated entries encompassing sequence, biological source, structural class, stereochemistry, terminal and chemical modifications, functional characterization, and hemolytic activity measurements;
- Cytotoxicity: cytotoxic peptide sequences were retrieved from Swiss-Prot/UniProt (Consortium, 2018) via advanced search filtering for sequence length between 2 and 50 residues and annotated with the cytotoxin keyword (KW-0800), yielding a set of experimentally validated cytotoxic peptides from diverse biological sources.
- Neurotoxicity: neurotoxic peptide sequences were retrieved from Swiss-prot/UniProt (Consortium, 2018) via advanced search filtering for sequence length between 2 and 50 residues and annotated with the neurotoxin keyword (KW-0528), yielding a set of experimentally validated neurotoxic peptides from diverse biological sources.

Together, these subtype datasets allowed us to evaluate whether NoiTox’s predictions align with distinct mechanistic categories of toxicity, providing a more biologically grounded assessment of model specificity beyond the general toxic/non-toxic dichotomy.

### Feature engineering

In the initial phase of this study, a comprehensive set of 8,461 sequence-derived descriptors was computed for each peptide using Biopython (Cock et al., 2009). The descriptor set included *Amino Acid Composition* (AAC; 20 features), *DiPeptide Composition* (DPC; 400 features), *TriPeptide Composition* (TPC; 8,000 features), *Composition of k-spaced Amino Acid Group Pairs* (CKSAAGP; 16 features), *Atomic Composition* (5 features), and several widely used *physicochemical properties* relevant to peptide toxicity, such as the Grand Average of Hydropathy (GRAVY) index, molecular mass, isoelectric point, hydrophobicity, and net charge (20 features), totalling 8,461 descriptors.

For the CKSAAGP, we calculated the frequency of pairs of amino acid groups separated by K residues in the peptide sequence. Amino acids were grouped based on their physicochemical properties, such as charge, polarity, and hydrophobic groups. This feature captures proximal interactions between amino acid groups that can significantly influence the peptide structure and function. Additionally, the CKSAAGP descriptors were normalized by the peptide length to ensure consistency across sequences of varying lengths.

Given the high dimensionality of the resulting feature space, an automated feature selection strategy was applied to identify the most informative descriptors. The Boruta algorithm (Kursa and Rudnicki, 2010), an all-relevant feature selection method based on ensemble tree models, was used to evaluate descriptor importance by comparing original features against randomized shadow features. This procedure classifies each variable as *confirmed, tentative*, or *rejected* based on its consistent contribution to predictive performance, reducing the risk of discarding relevant descriptors. This same descriptor computation and feature selection strategy was previously employed in the development of a cell-penetrating peptide prediction model (Bernardes-Loch et al., 2025), demonstrating its effectiveness in high-dimensional biological sequence classification tasks. Ultimately, 44 descriptors were confirmed as informative by Boruta and retained as the final feature subset used for model development, resulting in the model referred to as NoiTox (Bo).

### Evaluation strategy

#### Model training

Three distinct architectures were explored during model training, spanning classical machine learning, deep learning, and graph-based deep learning paradigms, providing a comprehensive comparison of approaches with fundamentally different inductive biases and representational strategies.

##### Classical machine learning (ML)

A set of supervised learning algorithms, including Extremely Randomized Trees (ERT), Random Forest (RF), XGBoost (XGB), Support Vector Machines (SVM), and Multilayer Perceptrons (MLP), was evaluated on the peptide toxicity training dataset. Two feature representations were initially compared during algorithm selection: the full 8,461-descriptor set reduced via Singular Value Decomposition (SVD), with performance assessed using up to 100 retained components, and the compact 44-descriptor subset identified by Boruta feature selection. Model robustness was examined through 10-fold cross-validation, with metrics averaged across folds. ERT coupled with Boruta-selected features achieved the highest Matthews Correlation Coefficient (MCC) across all evaluated combinations and was therefore selected as the final classical machine learning model, referred to as *NoiToxML (Bo)*. For comparison, we additionally report *NoiToxML (full)*, an ERT classifier trained on the complete, non-reduced 8,461-descriptor set (without SVD), using the same hyperparameters as NoiToxML (Bo).

##### Deep learning (DL)

Each peptide sequence was encoded using the ESM-2 protein language model, which generates contextual embeddings from evolutionary and biochemical information learned across hundreds of millions of protein sequences. A fixed-length representation was obtained by averaging per-residue embeddings from the final layer, a computationally efficient strategy widely adopted in the protein language model literature for variable-length sequences, which captures both local and global sequence properties without manual feature engineering. These vectors served as input to an MLP classifier trained with binary cross-entropy loss and the Adam optimizer, resulting in the model referred to as *NoiToxDP*.

##### Graph-based deep learning (GNN)

ESM-2 per-residue embeddings served as node features in a fully connected residue graph. Given that no structural data is required, a fully connected graph was adopted to capture all potential residue–residue interactions from sequence alone, enabling the model to learn long-range dependencies without relying on experimentally determined or predicted structures. Graph convolution was performed through three GATv2 layers, with learned attention pooling aggregating node embeddings into a graph-level representation. In addition, the 44 Boruta-selected descriptors were provided as graph-level global features, projected through a linear layer (with layer normalization and ELU activation) to 64 dimensions and concatenated with the pooled graph representation before the binary classification head. Training used binary cross-entropy loss with early stopping based on the validation loss, resulting in the model referred to as *NoiToxGP*.

##### Model parameters and evaluation

Full hyperparameter details for NoiToxML (Bo), NoiToxML (full), NoiToxDP, and NoiToxGP are provided in the Supplementary Material (section *Model parameters and evaluation*, Table S1). The 44 Boruta-selected descriptors, used as input by NoiToxML (Bo) and as global graph-level features by NoiToxGP, are listed in Supplementary Table S2. The tree ensembles were trained on the full training set, whereas for NoiToxDP and NoiToxGP 10% of the training set (stratified by class) was held out for early stopping. The random seed controls the tree ensembles, this validation split, weight initialization and data shuffling. All models were trained and evaluated with five random seeds (42–46), and the NoiTox results in the main text are means over these five runs; standard deviations are reported in Supplementary Tables S3–S4.

#### Comparison with alternative methods

NoiTox was evaluated and benchmarked against state-of-the-art approaches using a consistent set of performance metrics, including Sensitivity (SN), Specificity (SP), Accuracy (ACC), and Matthews Correlation Coefficient (MCC). Two evaluation strategies were used depending on baseline availability. For ToxiPep, PepNet, and both variants of ToxinPred2.0 (RF and Hybrid), the officially released pre-trained models were executed locally (ToxiPep, ToxinPred2.0) or via the method’s public webserver (PepNet), without retraining, on the test and independent datasets originally distributed for ToxiPep. The resulting metrics closely matched those reported in the original publications: for ToxinPred2.0, a decision-threshold sweep (0.1– 2.0, covering the full range of both the RF probability score and the wider Hybrid Score) confirmed that the default threshold (0.5) reproduces the published comparison most closely for both variants (Supplementary section *NoiTox results compared with state-of-the-art methods* Tables S5 and S6). The sequence-level outputs from these four re-executions enabled the misclassification overlap analysis described in the Incorrect predictions section and paired significance testing across all models sharing sequence-level predictions. For CSM-Toxin specifically, local re-execution was attempted both via its public API and its webserver interface; neither completed within a practical timeframe, so sequence-level predictions from this method could not be obtained, and CSM-Toxin was consequently excluded from the paired significance testing described below.

## Results and discussion

This section presents the experimental evaluation of NoiTox across three complementary model architectures. We first report the comparative assessment of classical machine learning algorithms that motivated the selection of ERT with Boruta-selected features as NoiToxML (Bo). We then benchmark the four NoiTox variants and their consensus against five established peptide toxicity predictors on standard benchmark datasets. Finally, we present a systematic analysis of sequences consistently misclassified across all models, cross-referencing them with hemolytic, cytotoxic, and neurotoxic databases to identify candidate label noise.

### NoiTox cross-validation analysis

As a first step, the classification performance of ERT, RF, XGB, SVM, and MLP was examined using an identical training dataset and feature matrix. All algorithms were assessed under a consistent 10-fold cross-validation strategy to ensure a fair comparison. Multiple dataset configurations were explored, preserving the same peptide sequences and descriptors while applying distinct preprocessing and modeling strategies.

Among the evaluated algorithms, Extremely Randomized Trees demonstrated superior predictive performance. In addition to its classification accuracy, this method enabled an effective assessment of feature relevance through the Boruta algorithm, which reduced the original 8,461-descriptor space to a compact subset of 44 confirmed informative descriptors. The resulting model, referred to as NoiToxML (Bo), is presented in Table 2.

**Table 2.**
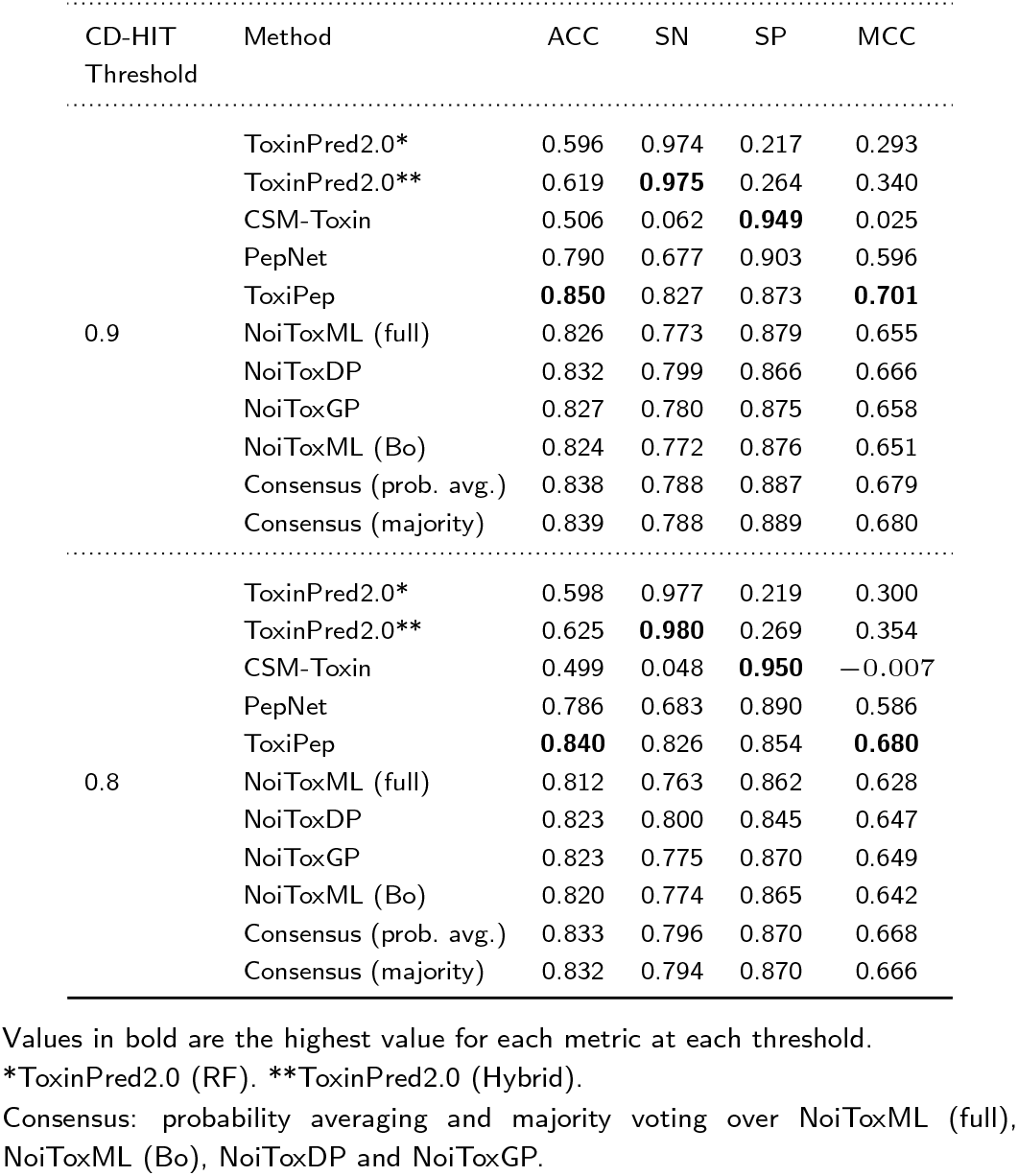
Comparison of NoiTox metrics with competing methods under different CD-HIT thresholds. NoiTox values are means over five random seeds; standard deviations in Supplementary section *Model parameters and evaluation* Tables S3–S4.

### NoiTox results compared with state-of-the-art methods

#### Benchmark test sets: CD-HIT 0.8 and CD-HIT 0.9

To ensure a fair comparison, the performance of NoiTox was evaluated against five established peptide toxicity predictors: ToxinPred2.0 (RF), ToxinPred2.0 (Hybrid), CSM-Toxin, PepNet, and ToxiPep. All methods were assessed using the validation datasets originally introduced by ToxiPep, which consist exclusively of experimentally validated natural peptides and are widely used as benchmarks in peptide toxicity prediction studies.

These datasets were clustered using two sequence similarity thresholds (0.8 and 0.9), resulting in sets of 1,388 and 1,766 peptides, respectively. Both datasets are fully balanced between toxic and non-toxic peptides (694/694 at the 0.8 threshold and 883/883 at the 0.9 threshold, as detailed in Table 1), allowing evaluation of model performance independent of class-distribution effects at two distinct levels of sequence redundancy reduction. The comparative results for both thresholds are summarized in Table 2.

Despite a more compact and interpretable design, the NoiTox variants remain competitive with ToxiPep across both thresholds, although ToxiPep retains the highest ACC and MCC on both CD-HIT test sets. At the 0.9 threshold, ToxiPep reaches ACC 0.850 and MCC 0.701, followed by the NoiTox consensus (ACC 0.838– 0.839, MCC 0.679–0.680), while the individual NoiTox variants reach MCC values of 0.651–0.666 (NoiToxDP being the best). At the 0.8 threshold, ToxiPep again leads (ACC 0.840, MCC 0.680), with the NoiTox consensus the closest configuration (MCC 0.666–0.668) and NoiToxGP and NoiToxDP the closest individual variants (MCC 0.649 and 0.647). NoiToxDP has the highest sensitivity among the individual NoiTox variants at both thresholds (0.799–0.800), and all NoiTox configurations avoid the specificity collapse of the sensitivity-biased ToxinPred2.0 variants. This shows that competitive toxicity prediction is attainable without ToxiPep’s multimodal complexity, though the best individual NoiTox variant differs by threshold. The narrow margins between top models, however, should be interpreted in light of benchmark labeling limitations, which we examine in subsection 3.5.

Two of the external methods warrant closer examination given their extreme SN/SP trade-offs: ToxinPred2.0 (evaluated in its RF and Hybrid variants) and CSM-Toxin. The very low specificity of ToxinPred2.0 at its default comparison threshold (SP *≈* 0.22–0.27 in Table 2) is explained by our threshold sweep (Supplementary Tables S5–S6): at threshold 0.5, sensitivity is near-maximal (SN *>* 0.97) at the cost of specificity for both variants, and specificity only rises to competitive levels (SP *>* 0.8) at thresholds of 0.8–0.9, where sensitivity falls substantially. This indicates a threshold miscalibration for this benchmark rather than a fundamental limitation of the underlying classifiers. Threshold-independent discrimination performance (AUROC/AUPRC, Supplementary Table S7) is consistent with this interpretation: on both CD-HIT test sets, ToxinPred2.0 reaches AUROC values of 0.81–0.85, well above what its MCC at the default threshold (0.29–0.35) would suggest, although lower than those of ToxiPep and the NoiTox models (0.89– 0.93). Its apparent weakness therefore stems mainly from threshold miscalibration, with a moderate ranking ability. Because we were unable to execute CSM-Toxin, either through its public API or its web server, we cannot independently diagnose the cause of its near-random performance (MCC *≈* 0 at CD-HIT 0.9, negative at CD-HIT 0.8); we report the values as published by (Guan et al., 2025).

#### Summary of comparison with state-of-the-art method

The comparative results obtained under both CD-HIT similarity thresholds demonstrate that NoiTox exhibits highly competitive and, in some cases, superior performance compared to established peptide toxicity predictors. At the 0.9 threshold, ToxiPep achieved the highest ACC and MCC (0.850 and 0.701), while the NoiTox consensus (ACC 0.838–0.839, MCC 0.679–0.680) and the individual NoiTox models (MCC 0.651–0.666) maintained strong and balanced performance, outperforming PepNet (MCC 0.596), CSM-Toxin, and both ToxinPred2.0 variants in ACC and MCC. At the 0.8 threshold, ToxiPep retained the highest ACC and MCC, with the NoiTox consensus the closest configuration (ACC 0.832–0.833, MCC 0.666–0.668) and NoiToxGP the closest individual variant (ACC 0.823, MCC 0.649), confirming balanced performance even at lower sequence similarity.

While ToxinPred2.0 models exhibited very high sensitivity, this behavior was accompanied by extremely low specificity, indicating a strong bias toward predicting peptides as toxic. Conversely, CSM-Toxin achieved high specificity at the cost of markedly reduced sensitivity, resulting in near-random or negative MCC values, particularly at the 0.8 threshold. In contrast, all NoiTox models maintained a well-balanced trade-off between sensitivity and specificity, leading to stable MCC values across both thresholds.

The consistent performance of NoiTox across datasets clustered at 0.8 and 0.9 similarity indicates robustness to redundancy reduction, while generalization beyond the validation distribution is demonstrated on the independent dataset (Table 3). NoiToxML (Bo) and NoiToxML (full) achieve this performance relying exclusively on interpretable physicochemical, compositional, and atomic descriptors, while NoiToxDP (embeddings only) and NoiToxGP (embeddings combined with the Boruta descriptors as global features) indicate that ESM-2 embeddings provide complementary representational power without sacrificing predictive balance.

Taken together, these results demonstrate that NoiTox offers a competitive alternative to state-of-the-art peptide toxicity predictors, approaching ToxiPep on the CD-HIT test sets and exceeding it on the independent dataset, while collectively covering the spectrum from interpretable feature-based classification to sequence-aware deep learning representations.

**Table 3.** Comparison of NoiTox metrics with competing method on the independent dataset. NoiTox values are means over five random seeds; standard deviations in Supplementary *Model parameters and evaluation* in Tables S3–S4.

| CD-HIT Threshold | Method | ACC | SN | SP | MCC |
| --- | --- | --- | --- | --- | --- |
| 0.9 | ToxiPep | 0.835 | 0.825 | 0.846 | 0.670 |
|  | NoiToxML (full) | 0.849 | 0.764 | **0.934** | 0.709 |
|  | NoiToxDP | 0.825 | **0.846** | 0.803 | 0.650 |
|  | NoiToxGP | 0.810 | 0.798 | 0.822 | 0.621 |
|  | NoiToxML (Bo) | **0.863** | 0.813 | 0.913 | **0.730** |
|  | Consensus (prob. avg.) | 0.861 | 0.833 | 0.890 | 0.724 |
|  | Consensus (majority) | **0.863** | 0.835 | 0.891 | 0.727 |
| 0.8 | ToxiPep | 0.815 | 0.821 | 0.810 | 0.631 |
|  | NoiToxML (full) | 0.851 | 0.782 | **0.919** | 0.708 |
|  | NoiToxDP | 0.818 | **0.859** | 0.777 | 0.638 |
|  | NoiToxGP | 0.808 | 0.793 | 0.822 | 0.619 |
|  | NoiToxML (Bo) | 0.850 | 0.812 | 0.887 | 0.702 |
|  | Consensus (prob. avg.) | **0.856** | 0.834 | 0.878 | **0.713** |
|  | Consensus (majority) | 0.855 | 0.833 | 0.878 | 0.712 |

Values in bold are the highest value for each metric at each threshold.
Consensus: probability averaging and majority voting over NoiToxML (full), NoiToxML (Bo), NoiToxDP and NoiToxGP.

To further assess model generalization, NoiTox models trained on the CD-HIT 0.8 and 0.9 datasets were evaluated on an independent competitive benchmark, with results compared against ToxiPep as the strongest baseline. The results are summarized in Table 3.

On the independent dataset, the NoiTox models generally surpassed ToxiPep in MCC. At CD-HIT 0.9, NoiToxML (Bo) (0.730), the probability-averaging and majority-voting consensus (0.724 and 0.727) and NoiToxML (full) (0.709) exceeded ToxiPep (0.670), whereas NoiToxDP (0.650) and NoiToxGP (0.621) trailed it. At CD-HIT 0.8, the consensus (0.713 and 0.712), NoiToxML (full) (0.708), NoiToxML (Bo) (0.702) and, marginally, NoiToxDP (0.638) exceeded ToxiPep (0.631), while NoiToxGP was lower (0.619). NoiToxML (full) attained the highest specificity at both thresholds (0.934 and 0.919) and NoiToxDP the highest sensitivity (0.846 and 0.859).

Across both thresholds and on both types of evaluation set, NoiToxDP exhibited higher sensitivity at the cost of reduced specificity relative to NoiToxML (Bo) (for example, SN 0.859 versus 0.812 and SP 0.777 versus 0.887 on the independent dataset at CD-HIT 0.8), reflecting the different inductive biases of sequence-embedding-based representations compared to physicochemical descriptors. NoiToxGP was comparable to NoiToxML (Bo) on the CD-HIT test sets and showed lower sensitivity and specificity on the independent dataset, with the largest variability across seeds. This complementarity suggests that ensemble or consensus strategies combining the three architectures could further improve predictive balance. However, preliminary voting experiments revealed that certain sequences are consistently misclassified across all architectures regardless of the aggregation strategy employed, indicating that these errors are not attributable to individual model limitations. To assess whether such sequences could simply be discarded as label noise, we removed from the training sets those that all four models contradicted in out-of-fold predictions, retrained all models, and evaluated them on unmodified test sets, using the random removal of the same number of sequences as a control. Cleaning did not yield a detectable improvement in the ranking performance of any model and, relative to the control, reduced the AUROC of NoiToxML and of the consensus in all evaluations (ΔAUROC ranging from *−*0.004 to *−*0.027), whereas NoiToxDP and NoiToxGP were essentially unaffected. These sequences therefore carry useful information and are better treated as candidates for curation than removed automatically (details in the Supplementary Material, section *Consensus-based cleaning of the training data* in Tables S13–S15). This pattern motivates a dedicated investigation into the nature of these persistently misclassified sequences, presented in the *Incorrect predictions* section.

We formalized this ensemble intuition by combining the four NoiTox variants through probability averaging (mean predicted probability, threshold 0.5) and majority voting, both implemented in the NoiTox.py orchestrator distributed with this work. Probability averaging gave a higher MCC than the best individual variant on the two CD-HIT test sets (by 0.019 at 0.8 and 0.013 at 0.9; Supplementary Table S12, Supplementary Material, section *Consensus strategies and recommended use*) and a comparable one, within seed-to-seed variability, on the independent dataset (0.713 versus 0.708 at 0.8; 0.724 versus 0.730 at 0.9). We recommend probability averaging as the default consensus strategy for end users.

### Benchmark validity: training–benchmark overlap and its effect on performance

Benchmark comparisons are only informative to the extent that training and evaluation data are genuinely independent. We therefore quantified sequence overlap between our training partitions and every benchmark set used in this study, the CD-HIT 0.8/0.9 test sets and the independent dataset, at two levels: *exact* matches (identical sequences) and *above-threshold* matches (sequences falling in the same CD-HIT cluster as a training sequence, at the same identity threshold used to construct that partition, i.e., 0.8 or 0.9).

#### Training-test overlap

No exact matches were found between either training partition and its corresponding test set. Above-threshold clustering identified a small number of matches: 27 of 1,388 test sequences (1.94%) at CD-HIT 0.8, and 29 of 1,766 (1.64%) at CD-HIT 0.9. Critically, every one of these matches was *cross-label* (a toxic test sequence clustering with a non-toxic training sequence, or vice versa); none matched a training sequence of the same label. This pattern is inconsistent with data leakage, which would be expected to inflate performance through same-label duplication, and is instead consistent with the label-noise-by-omission phenomenon developed in the Section Incorrect predictions: sequences that are highly similar in composition receive conflicting toxicity labels because negative labels are assigned by the absence of annotation rather than by experimental evidence. No sequences were therefore removed from the CD-HIT 0.8/0.9 test sets, and the metrics reported in Table 2 are unaffected by this analysis.

#### Training-independent overlap

The independent dataset, by contrast, showed substantial overlap with the training partitions. Exact matches were rare (2 of 958 sequences against the CD-HIT 0.8 training set; 0 against CD-HIT 0.9), but both exact matches involved a direct label contradiction: the sequences `GILDTIKSIASKVWNSKTVQDLKRKGINWVANKLGVSPQAA and FLPVVAGLAAKVLPSIICAVTKKC`, labeled toxic in the independent set, appear verbatim as non-toxic entries in the CD-HIT 0.8 training partition.

Above-threshold overlap was far more extensive: 352 of 958 independent sequences (36.74%) clustered with a CD-HIT 0.8 training sequence, and 287 of 958 (29.96%) with a CD-HIT 0.9 training sequence. Unlike the training-test comparison (Table 4), this overlap was predominantly *same-label* : 80.40% of the affected sequences (CD-HIT 0.8) and 88.50% (CD-HIT 0.9) matched a training sequence sharing the same toxic/non-toxic annotation, with a further 12.78% and 8.01%, respectively, matching training sequences of both labels simultaneously. Only 6.82% and 3.48% matched exclusively cross-label training sequences. This same-label-dominated pattern is the signature of genuine data leakage: models may have learned to recognize near-duplicate compositional profiles rather than generalizing to unseen toxic determinants.

**Table 4.** Label composition of above-threshold sequence overlap between training partitions and each benchmark set. A pair is counted for every individual training sequence sharing a CD-HIT cluster with a benchmark sequence (a benchmark sequence with multiple training matches contributes multiple pairs). pos-pos and neg-neg indicate same-label overlap (a signature of data leakage); cross indicates a toxic sequence clustering with a non-toxic sequence or vice versa (a signature of label noise by omission, not leakage).

| Benchmark | CD-HIT | Pairs | pos-pos | neg-neg | cross |
| --- | --- | --- | --- | --- | --- |
| Test | 0.8 | 34 | 0 (0%) | 0 (0%) | 34 (100%) |
| Test | 0.9 | 30 | 0 (0%) | 0 (0%) | 30 (100%) |
| Independent | 0.8 | 427 | 175 (41.0%) | 167 (39.1%) | 85 (19.9%) |
| Independent | 0.9 | 313 | 131 (41.9%) | 148 (47.3%) | 34 (10.9%) |

#### Performance after removing overlapping sequences

To assess the impact of this leakage, we removed every independent-dataset sequence with at least one same-label match in the corresponding training partition (328 sequences for CD-HIT 0.8, leaving 630; 277 sequences for CD-HIT 0.9, leaving 681) and re-evaluated all previously trained models, without retraining, on the reduced sets. Sequences whose only overlap was cross-label were retained, consistent with their interpretation as label noise rather than leakage. Results are summarized in Table 5.

**Table 5.** Performance on the independent dataset before and after removing sequences with same-label overlap against the corresponding training partition. NoiTox values are means over five random seeds.

| CD-HIT | Method | ACC | SN | SP | MCC |
| --- | --- | --- | --- | --- | --- |
| 0.9, before | ToxiPep | 0.835 | 0.825 | 0.846 | 0.670 |
|  | NoiToxML (full) | 0.849 | 0.764 | 0.934 | 0.709 |
|  | NoiToxDP | 0.825 | 0.846 | 0.803 | 0.650 |
|  | NoiToxGP | 0.810 | 0.798 | 0.822 | 0.621 |
|  | NoiToxML (Bo) | 0.863 | 0.813 | 0.913 | 0.730 |
| 0.9, after | ToxiPep | 0.832 | 0.797 | 0.870 | 0.668 |
|  | NoiToxML (full) | 0.740 | 0.533 | 0.958 | 0.539 |
|  | NoiToxDP | **0.853** | 0.834 | 0.874 | 0.707 |
|  | NoiToxGP | 0.837 | **0.900** | 0.771 | 0.678 |
|  | NoiToxML (Bo) | 0.718 | 0.490 | **0.958** | 0.503 |
| 0.8, before | ToxiPep | 0.815 | 0.820 | 0.810 | 0.631 |
|  | NoiToxML (full) | 0.851 | 0.782 | 0.919 | 0.708 |
|  | NoiToxDP | 0.818 | 0.859 | 0.777 | 0.638 |
|  | NoiToxGP | 0.808 | 0.793 | 0.822 | 0.619 |
|  | NoiToxML (Bo) | 0.850 | 0.812 | 0.887 | 0.702 |
| 0.8, after | ToxiPep | 0.836 | 0.777 | 0.892 | 0.675 |
|  | NoiToxML (full) | 0.767 | 0.538 | 0.982 | 0.585 |
|  | NoiToxDP | 0.808 | 0.816 | 0.800 | 0.616 |
|  | NoiToxGP | **0.829** | 0.813 | 0.843 | **0.657** |
|  | NoiToxML (Bo) | 0.743 | 0.485 | **0.985** | 0.548 |

The effect of removing overlapping sequences is markedly uneven across architectures. ToxiPep, NoiToxDP, and NoiToxGP show stable or slightly improved MCC after removal (e.g., NoiToxDP: 0.650 to 0.707 at CD-HIT 0.9; NoiToxGP: 0.621 to 0.678), indicating that their performance on the independent dataset reflected genuine generalization rather than memorized near-duplicates. In sharp contrast, both NoiToxML variants collapse: MCC for NoiToxML (Bo) falls from 0.730 to 0.503 (CD-HIT 0.9) and from 0.702 to 0.548 (CD-HIT 0.8), driven almost entirely by a loss of sensitivity (0.813 to 0.490 and 0.812 to 0.485, respectively) while specificity rises to near-ceiling (0.958–0.985). This pattern, a selective loss of sensitivity concentrated in the removed subset, with specificity unaffected or improved, is the expected signature of a model that had partially memorized compositional profiles of near-duplicate positive sequences rather than learning generalizable toxicity determinants. We attribute this susceptibility to the reliance of NoiToxML on global physicochemical and compositional descriptors, which vary little between near-identical sequences, whereas the learned representations underlying NoiToxDP and NoiToxGP (ESM-2 embeddings, with or without residue-level message passing over a fully connected graph) appear more robust to this specific failure mode.

These results directly qualify the claims made in the Section NoiTox results compared with state-of-the-art methods: after correcting for training–independent overlap, NoiToxML (Bo) is no longer the strongest-performing NoiTox variant on the independent dataset, and its apparent superiority over ToxiPep does not survive the removal of same-label overlapping sequences.

#### Statistical significance of comparative claims

To determine which of the MCC differences reported above are statistically robust, we compared every pair of models with sequence-level predictions (ToxiPep, PepNet, the four NoiTox variants and the probability-averaging consensus) on the CD-HIT 0.8 and 0.9 test sets and on the independent dataset (evaluated with the models trained at each CD-HIT threshold; PepNet was not evaluated on the independent dataset). For each of the five random seeds we applied McNemar’s test to the paired per-sequence correctness and computed a bootstrap 95% confidence interval (10,000 resamples) for the difference in MCC (ΔMCC). ToxiPep and PepNet are single evaluations, so variability across seeds comes only from the NoiTox models. We report the mean ΔMCC, the number of seeds in which each comparison was significant and, to account for the 15–21 comparisons per benchmark, Holm-adjusted *p*-values computed on the median McNemar *p* across seeds. Full results are reported in Supplementary Table S8.

On the CD-HIT test sets, ToxiPep had a significantly higher MCC than PepNet and than NoiToxML (full) at both thresholds (all five seeds; ΔMCC = 0.052 at 0.8 and 0.046 at 0.9; Holm *p* = 0.046 and 0.019) and than NoiToxML (Bo) at 0.9 (ΔMCC = 0.049; five of five seeds; Holm *p* = 0.009). Its differences from NoiToxDP, NoiToxGP (0.031–0.043) and NoiToxML (Bo) at 0.8 (0.038) were significant in two to five of the five seeds but did not remain significant after correction (Holm *p ≥* 0.09), and its difference from the NoiTox consensus was not significant (ΔMCC = 0.012 at 0.8 and 0.022 at 0.9; zero and one of five seeds). PepNet had a lower MCC than every NoiTox configuration (0.042–0.083), significantly so for the consensus at both thresholds and for all variants at 0.9 (Holm *p ≤* 0.036). Among the NoiTox variants, no two individual models differed significantly in more than one seed, whereas the consensus exceeded NoiToxML (full) in all five seeds at 0.8 and in three at 0.9, although not after correction.

On the independent dataset at CD-HIT 0.8, the NoiTox consensus had a significantly higher MCC than ToxiPep (ΔMCC = 0.083; five of five seeds; Holm *p* = 0.004), and NoiToxML (full) and (Bo) were also higher in all five seeds (0.077 and 0.071), but these two differences did not remain significant after correction (Holm *p* = 0.072). NoiToxDP and NoiToxGP did not differ from ToxiPep (zero and one of five seeds). At CD-HIT 0.9, NoiToxML (Bo) (0.060; five of five seeds) and the consensus (0.054; three of five) exceeded ToxiPep, NoiToxML (full) did so inconsistently (McNemar test significant in no seed, bootstrap in two), and no comparison was significant after correction. Within the NoiTox framework, the NoiToxML variants and the consensus were higher than NoiToxDP and NoiToxGP in most seeds (ΔMCC 0.059–0.109), NoiToxML (full), NoiToxML (Bo) and the consensus did not differ from one another, and NoiToxDP did not differ from NoiToxGP.

These results temper several comparisons made above. On the CD-HIT test sets ToxiPep is significantly better than the NoiToxML variants but statistically indistinguishable from the NoiTox consensus, the closest configuration. On the independent dataset, the NoiToxML variants and the consensus exceeded ToxiPep in most seeds, but after correction only the consensus at CD-HIT 0.8 remained significant, and the advantage of the NoiToxML variants was not maintained after removing sequences similar to the training data (previous section). NoiToxDP and NoiToxGP were statistically comparable to ToxiPep on the independent dataset but lower than the NoiToxML variants and the consensus. The evidence supports competitive performance, with the consensus statistically tied with ToxiPep on the CD-HIT test sets and above it on the independent dataset at CD-HIT 0.8.

The lower MCC of NoiToxML (full) after overlap correction is best interpreted by distinguishing two kinds of metrics. Accuracy, sensitivity, specificity and MCC depend on a fixed decision threshold (0.5 on the predicted probability), so they change when the model’s scores shift relative to that cutoff, even if the model still ranks toxic peptides above non-toxic ones. AUROC and AUPRC, in contrast, do not depend on any threshold and measure only that ranking. On the complete independent dataset, NoiToxML (full) ranked sequences well: its AUROC was the highest among the individual methods at both thresholds (0.926 and 0.934, versus 0.889 and 0.886 for ToxiPep; Supplementary Table S7). Its drop in MCC after overlap removal was driven by lower sensitivity (0.533–0.538) with specificity of 0.958– 0.982, which is compatible with a shift of its scores relative to the 0.5 threshold rather than with a loss of ranking ability, as we also observed for ToxinPred2.0 above.

### Feature importance

Interpretability remains a key challenge in peptide toxicity prediction, as most state-of-the-art approaches rely on high-dimensional embeddings that obscure the molecular determinants of toxicity. In contrast, NoiToxML (Bo) explicitly models physicochemical, compositional, and atomic-level properties, achieving competitive predictive performance while providing direct insight into the features associated with toxic and non-toxic sequences.

To investigate the contribution of individual descriptors, feature importance was analyzed using the Extremely Randomized Trees algorithm. The 20 most informative descriptors among the 44 Boruta-confirmed features are summarized in Figure 1, providing a focused view of the most discriminative variables without visual overload.

**Figure 1.**
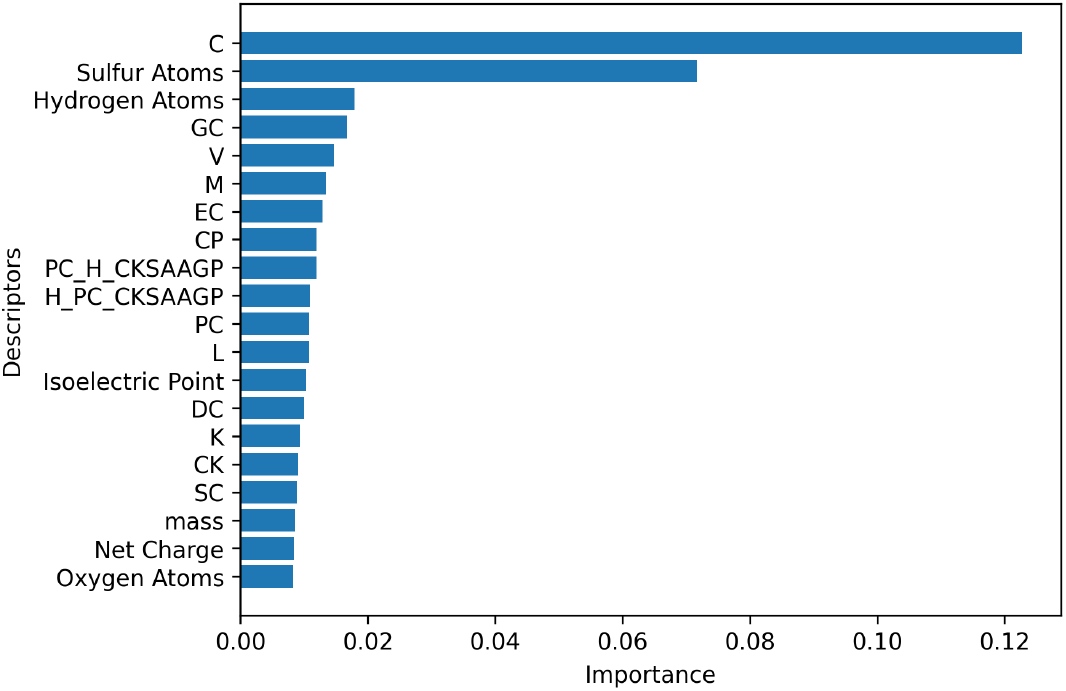
Top 20 independent descriptors for toxin classification identified by Extremely Randomized Trees algorithm. The abbreviations PC, and H represent the amino acid groups positively charged, and hydrophobic, respectively.

As shown in Table 2, NoiToxML (Bo) offers a substantially more compact representation than NoiToxML (full), relying on 44 rather than 8,461 descriptors, while achieving comparable performance on the CD-HIT test sets (MCC 0.651 versus 0.655 at 0.9; 0.642 versus 0.628 at 0.8) and on the independent dataset (0.730 versus 0.709 at 0.9; 0.702 versus 0.708 at 0.8; Table 3). NoiToxML (Bo) therefore combines a compact, interpretable representation, detailed below, with discriminative performance comparable to that of the embedding-based variants.

The feature importance analysis (Figure 1) indicates that descriptors related to cysteine (C) and sulfur atoms contribute substantially to peptide toxicity prediction. This observation is consistent with the chemical properties of cysteine, whose thiol (–SH) group is highly redox-active and participates in disulfide bond formation and other covalent interactions that influence peptide stability and biological activity (Schöneich, 2017). Experimental studies also indicate that excessive cysteine exposure can induce cellular damage, reflecting the need for tight regulation of its chemical reactivity (Shibui et al., 2017). In addition, sulfur atoms and reactive sulfur species can interact with cellular components involved in oxidative stress and toxicity pathways (Shinkai and Kumagai, 2019). Consistent with this interpretation, the comparison of descriptor distributions between toxic and non-toxic peptides further supports the relevance of cysteine- and sulfur-related features. As shown in Figure 2, toxic peptides present a higher normalized proportion of sulfur atoms and cysteine (C) than non-toxic sequences.

**Figure 2.**
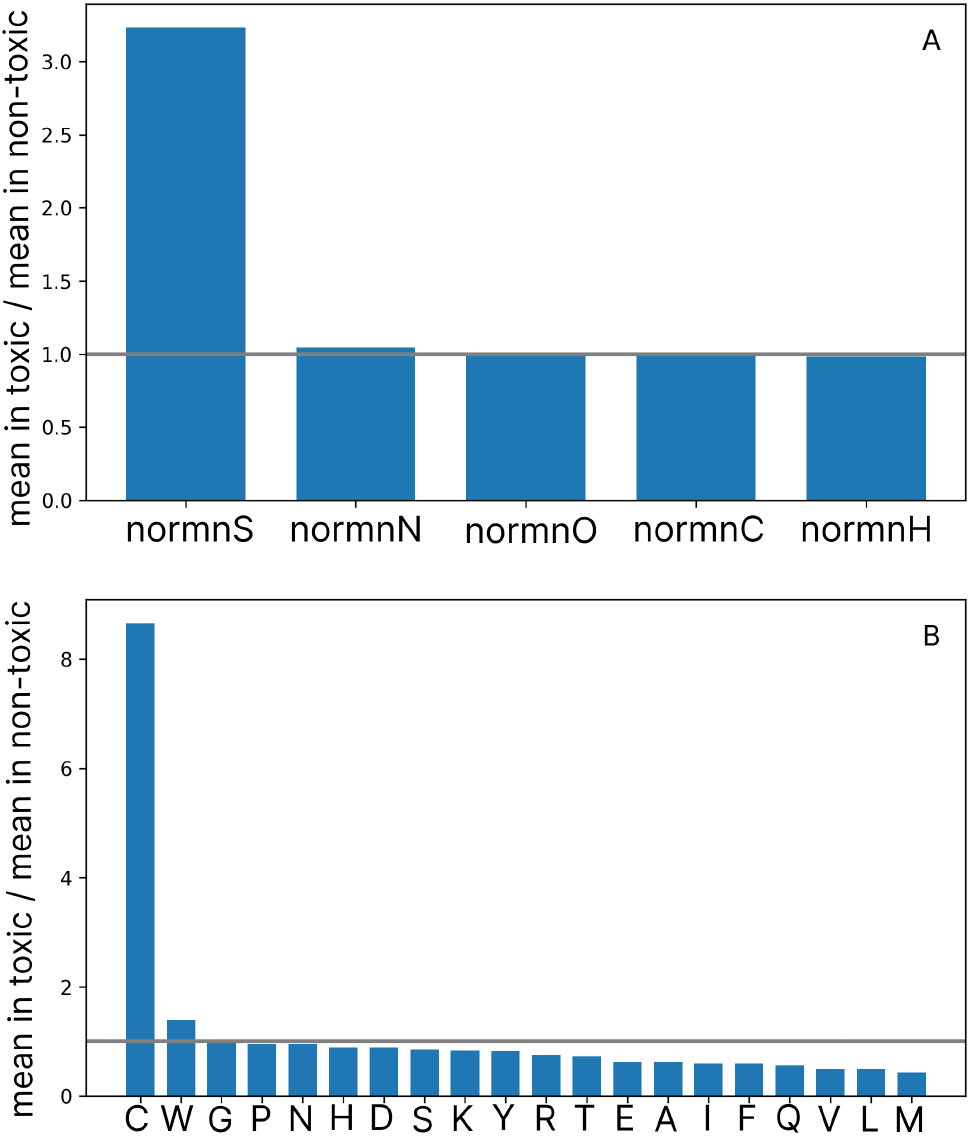
Mean ratio of normalized atomic (A) and amino acid (B) descriptors between toxic and non-toxic peptide groups. Each bar represents the ratio (mean in toxic / mean in non-toxic) of a specific feature. Values greater than 1 indicate a higher mean in the toxic group, whereas values less than 1 indicate a higher mean in the non-toxic group.

At the amino acid level, cysteine is enriched in toxic peptides, a pattern also reflected in residue-pair analysis (Figure 3), where several cysteine-containing dipeptides are overrepresented in the toxic class. These results indicate that the elevated importance of cysteine and sulfur descriptors reflects systematic compositional differences between toxic and non-toxic peptides rather than artifacts of feature selection.

**Figure 3.**
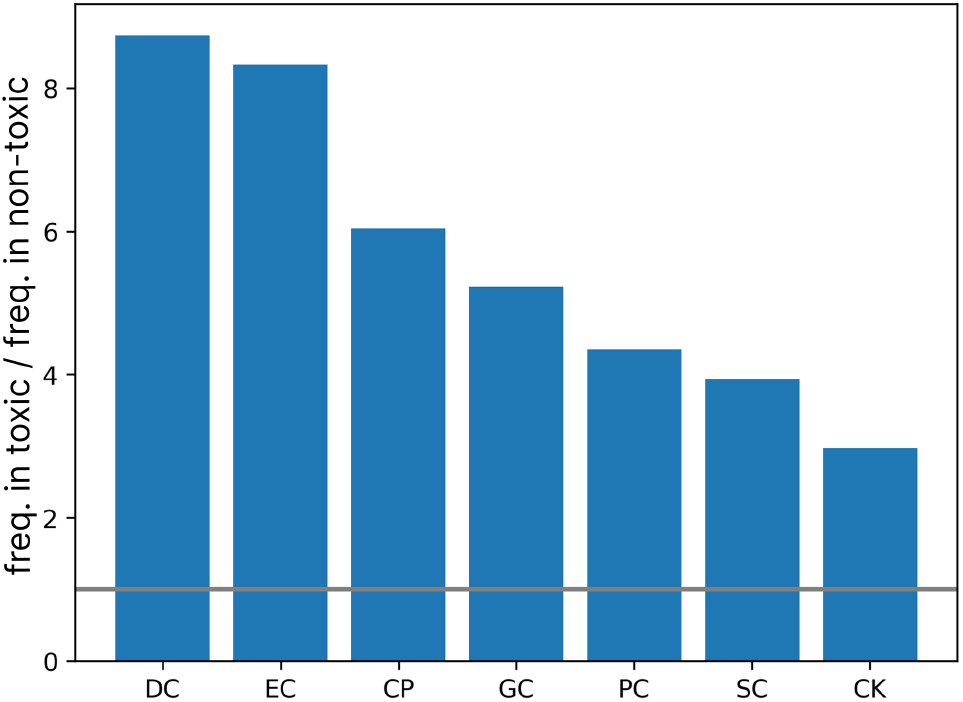
Frequency ratio of cysteine-containing dipeptide descriptors between toxic and non-toxic peptide groups. Each bar represents the ratio (frequency in toxic / frequency in non-toxic) of a specific descriptor. Values greater than 1 indicate higher frequency in the toxic group, whereas values less than 1 indicate higher frequency in the non-toxic group.

In contrast, methionine (M), although also sulfur-containing, shows the opposite distribution pattern and is more abundant in non-toxic peptides. This difference is consistent with its thioether side chain, which is less chemically reactive than the cysteine thiol group (Schöneich, 2017). Similarly, the enrichment of valine (V) and leucine (L) in non-toxic peptides agrees with previous studies indicating that aliphatic hydrophobic residues without reactive side chains often contribute to structural stability without inducing membrane disruption or hemolysis (Robles-Loaiza et al., 2022). These observations suggest that methionine, valine, and leucine may act as compositional indicators of non-toxic peptides.

### Incorrect predictions

In total, the models produced 643 unique misclassifications, including 490 in the test dataset (212 non-toxic and 278 toxic sequences, after removing duplicates shared between the 0.8 and 0.9 CD-HIT thresholds) and 153 in the independent dataset. A detailed summary of incorrect predictions across datasets and classes is presented in Table 6.

**Table 6.** Incorrect predictions in NoiTox. Freq. is the number of sequences in toxic and non-toxic subsets, CP is correct predictions and IP is incorrect predictions.

| CD-HIT Threshold | Label | Freq. | CP | IP |
| --- | --- | --- | --- | --- |
| 0.9 | Toxic | 883 | 724 | 159 |
|  | Non-toxic | 883 | 749 | 134 |
| 0.8 | Toxic | 694 | 549 | 145 |
|  | Non-toxic | 694 | 591 | 103 |
| Indep. | Toxic | 479 | 387 | 92 |
|  | Non-toxic | 479 | 418 | 61 |

To investigate the origin of these errors, we examined the distributions of selected descriptors for correctly and incorrectly predicted peptides. As illustrated in Figure 4, toxic peptides misclassified as non-toxic show Positively Charged Hydrophobic CKSAAGP values similar to those of correctly classified non-toxic peptides. A comparable trend is observed for Net Charge: toxic peptides typically display positive median values, whereas correctly classified non-toxic peptides are centered near zero (near-neutral medians). Non-toxic peptides misclassified as toxic shift toward positive net charge values, resembling the toxic distribution. These results indicate that similarities in CKSAAGP and Net Charge descriptors contribute to overlapping regions in the feature space where toxic and non-toxic peptides become difficult to distinguish.

**Figure 4.**
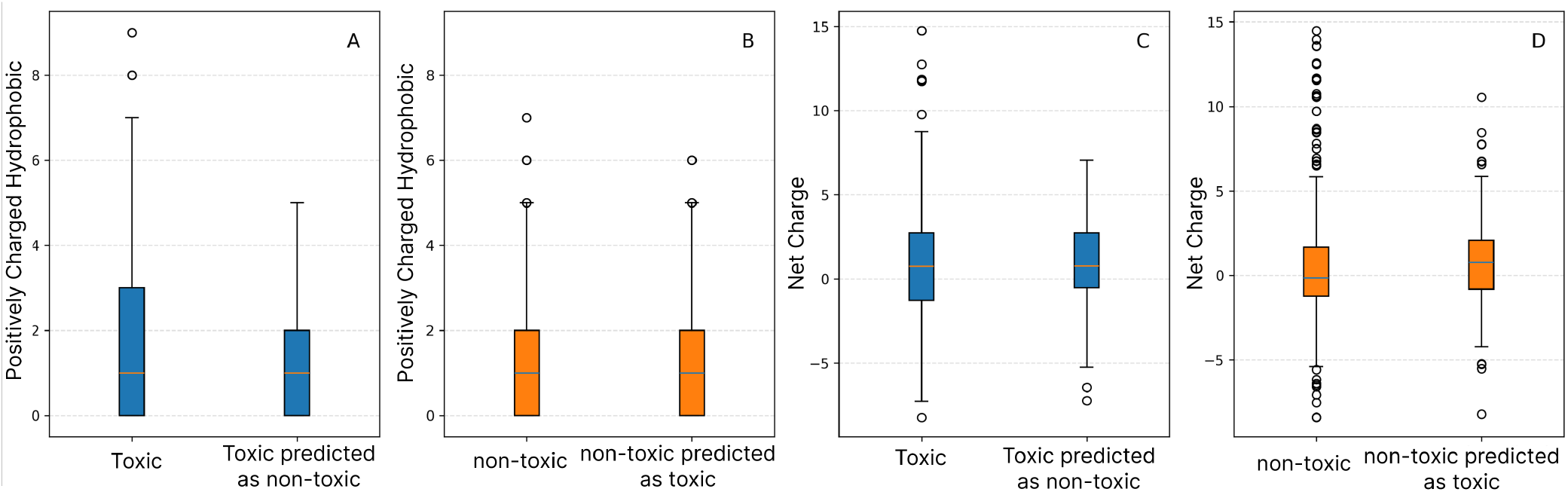
Distribution of Positively Charged Hydrophobic CKSAAGP and Net Charge descriptors for training and misclassified sequences. **A** Distribution of Positively Charged Hydrophobic CKSAAGP for positive training sequences and misclassified sequences. **B** Positively Charged Hydrophobic CKSAAGP distribution for negative training sequences and misclassified sequences. **C** Net charge distribution for positive training sequences and misclassified sequences. **D** Net charge distribution for negative training sequences and misclassified sequences.

A complementary z-score analysis (Figure 5) further clarifies the behavior of misclassified peptides. Toxic peptides predicted as non-toxic show minimal deviation from the correctly predicted non-toxic group, indicating strong similarity to the non-toxic feature profile captured by the model. In contrast, non-toxic peptides predicted as toxic show higher levels of cysteine and sulfur-related descriptors and lower levels of methionine and valine relative to the non-toxic reference group. These shifts move the peptides toward the toxic region of the feature space.

**Figure 5.**
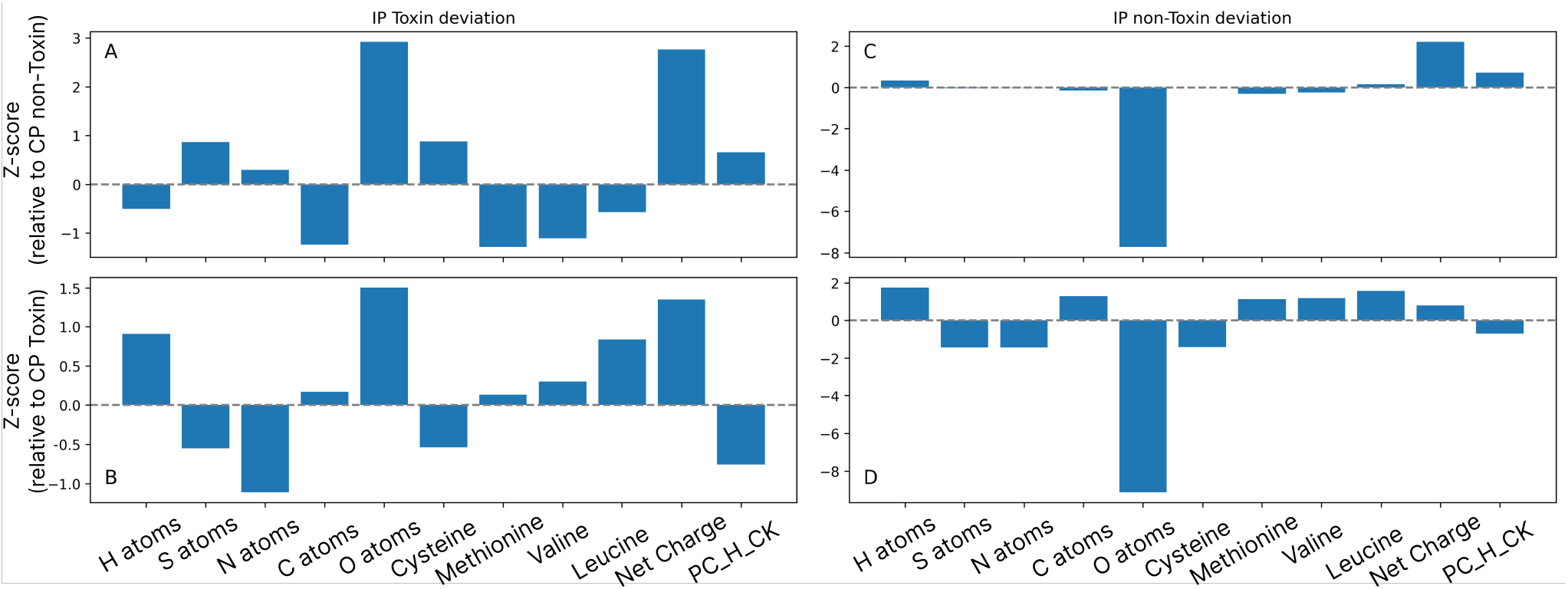
Z-score deviation of selected features for incorrectly predicted peptides relative to correctly predicted peptides. Bars represent feature-wise z-score deviations, where positive values indicate higher levels and negative values indicate lower levels relative to the reference group. **A** Incorrectly predicted toxic peptides relative to correctly predicted non-toxic peptides. **B** Incorrectly predicted toxic peptides relative to correctly predicted toxic peptides. **C** Incorrectly predicted non-toxic peptides relative to correctly predicted non-toxic peptides. **D** Incorrectly predicted non-toxic peptides relative to correctly predicted toxic peptides.

Overall, these results suggest that incorrect predictions arise primarily from overlapping compositional and physicochemical profiles between toxic and non-toxic peptides rather than from isolated anomalies in individual descriptors.

A complementary analysis revealed a structural limitation in the datasets used for training, testing, and independent validation. A search of sequences labeled as non-toxic against experimentally validated hemolytic, cytotoxic, and neurotoxic peptide databases identified inconsistencies across all evaluated partitions. Sequences were matched by exact string comparison of their amino acid sequences: a sequence was counted only when it was identical to an entry of the external database, and no similarity or identity threshold was applied. Near-identical or homologous sequences were therefore not counted, and the numbers reported below should be read as a lower bound of the sequences with conflicting annotation. The number of unique sequences carrying a negative label that matched at least one experimentally confirmed toxic entry is summarized in Table 7.

**Table 7.** Number of unique sequences labeled as non-toxic that matched experimentally validated toxic peptides across external databases, stratified by dataset partition and CD-HIT clustering threshold.

| Partition | Training set | Test set |
| --- | --- | --- |
| CD-HIT 0.9 | 37 (0.53%) | 08 (0.91%) |
| CD-HIT 0.8 | 26 (0.47%) | 10 (1.44%) |
| CD-HIT 0.9 + 0.8 (union) | 42 (0.51%) | 16 (2.30%) |
| Independent dataset | 9 (0.94%) |  |

A breakdown by toxicity subtype and database of origin is presented in Table 8. Across both clustering thresholds, the most prevalent sources of mislabeled sequences were neurotoxic peptides (UniProt/KW-0528), and hemolytic peptides (Hemolytic2), with cytotoxic and DBAASP-derived entries contributing to a lesser extent. In the independent validation set, the 9 affected sequences were distributed across cytotoxic (n = 5), hemolytic (n = 1), and neurotoxic/UniProt (n = 3) categories.

**Table 8.** Distribution of label-noise sequences by toxicity subtype and source database, stratified by CD-HIT threshold and dataset partition.

| CD-HIT | Source database | Training | Test |
| --- | --- | --- | --- |
| 0.8 | Cytotoxic (general) | 4 | 1 |
|  | Cytotoxic (DBAASP) | 2 | 0 |
|  | Hemolytic (Hemolytic2) | 10 | 3 |
|  | Neurotoxic (UniProt) | 10 | 6 |
| 0.9 | Cytotoxic (general) | 7 | 1 |
|  | Cytotoxic (DBAASP) | 0 | 2 |
|  | Hemolytic (Hemolytic2) | 15 | 1 |
|  | Neurotoxic (UniProt) | 15 | 4 |
| *Independent dataset (no threshold subdivision):* |  |  |  |
|  | Cytotoxic | 5 |  |
|  | Hemolytic (Hemolytic2) | 1 |  |
|  | Neurotoxic (UniProt) | 3 |  |

To test whether this pattern reflects a systematic signal rather than chance co-occurrence, we compared the rate at which negative-labeled sequences matched an experimentally validated toxicity database between two groups, per model and per partition: sequences misclassified as toxic (false positives) versus sequences correctly classified as non-toxic (true negatives). Fisher’s exact test (one-sided, testing for an elevated match rate among misclassified sequences) confirmed a significant enrichment in 6 of 8 model-partition combinations (Supplementary Table S9), with odds ratios ranging from 6.5 to 48.2 among significant results (*p <* 0.05). The two non-significant comparisons (NoiToxML, CD-HIT 0.8) involved small absolute counts (one matched sequence in each misclassified group) and are likely underpowered rather than indicating an absent effect, given that the same models showed strong, significant enrichment at CD-HIT 0.9 (*p <* 0.001 and *p* = 0.026, respectively). These results indicate that misclassified negative-labeled sequences are enriched for sequences annotated as toxic in external databases, rather than randomly distributed with respect to this independent evidence.

This finding suggests that the negative set, originally constructed through UniProt filters based on the absence of explicit toxicity annotation, may contain functionally toxic peptides that were simply not yet annotated as such. This phenomenon, known as *label noise by omission*, is well documented in bioinformatics classification tasks where the negative class is defined by the absence of evidence rather than evidence of absence. Critically, this issue is not specific to peptide toxicity prediction. Sidorczuk et al. (Sidorczuk et al., 2022) demonstrated, across 660 predictive models and 11 negative sampling strategies, that benchmark performance in antimicrobial peptide prediction is systematically determined by the choice of negative data: models trained and evaluated on datasets sharing the same sampling method exhibit artificially inflated and non-reproducible performance estimates. Taken together, these findings suggest that a fraction of the persistent misclassifications observed in NoiTox may reflect genuine toxic activity in sequences incorrectly assigned to the negative class, rather than limitations of the models themselves.

This issue becomes particularly relevant when one observes that all three NoiTox models, NoiToxML (Bo), NoiToxDP, and NoiToxGP, together with ToxiPep, used here as an external reference applied to the same sequence set, consistently commit errors on the same sequences, a behavior illustrated in the error overlap diagram (Figure 6). If the models were simply underfitted or incapable of capturing toxic function, a more heterogeneous distribution of errors across distinct architectures would be expected. The convergence of errors points, instead, to a contradictory signal in the data itself: the sequences in question carry characteristics that all models consistently associate with toxicity, yet their original label classifies them as non-toxic. The 20 sequences are in the supplementary material in Table S10.

**Figure 6.**
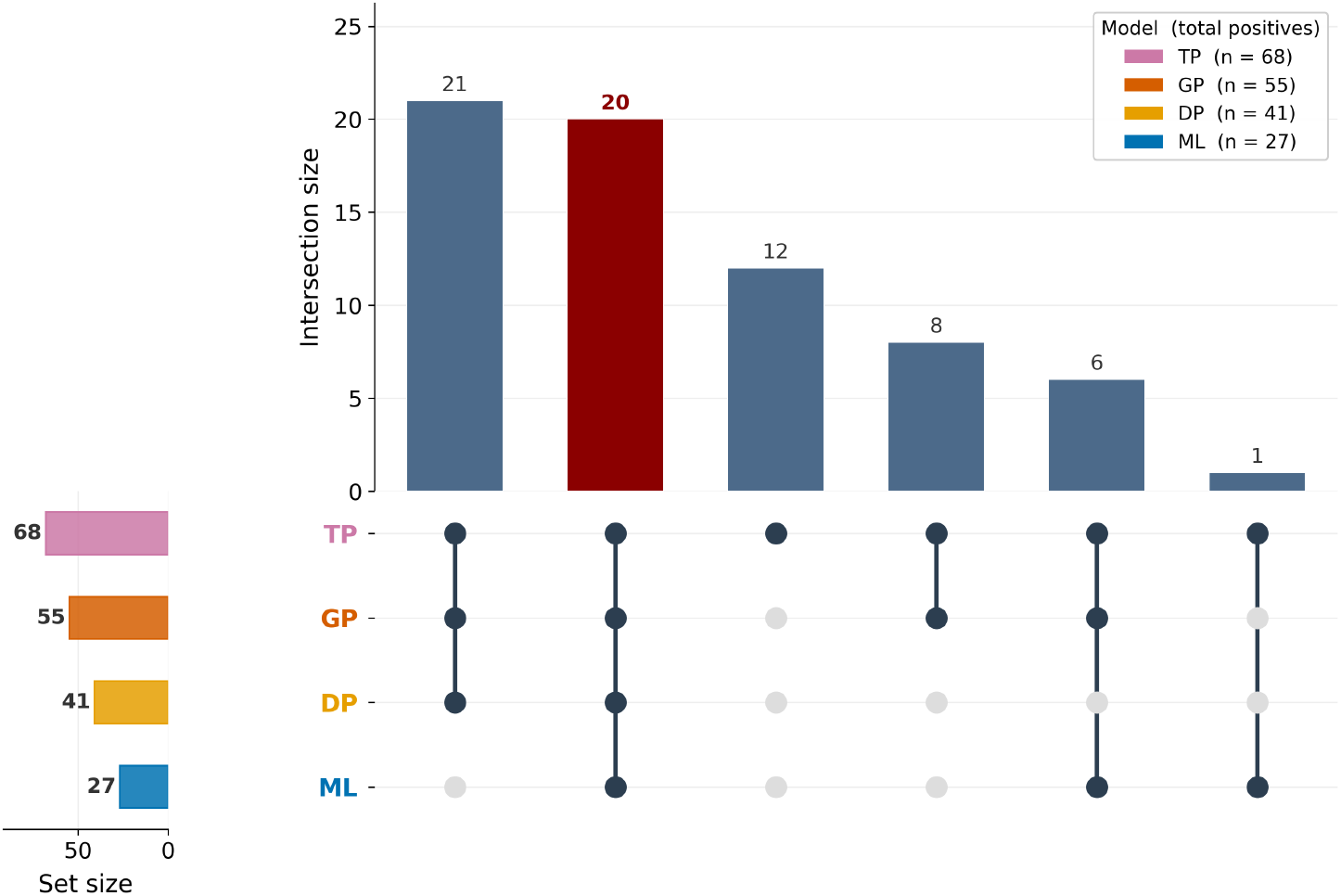
Consensus analysis of positive predictions across four computational models for toxic peptide identification. UpSet plot displaying the intersection pattern of non-toxic peptide sequences predicted as toxic by four independent models: NoiToxML (Bo) (ML; n = 27), NoiToxDP (DP; n = 41), NoiToxGP (GP; n = 55), and ToxiPep (TP), used as an external reference (n = 68) . Bars represent the number of sequences exclusively shared by each combination of models, sorted by decreasing intersection size. The dark red bar (n = 20) indicates sequences unanimously predicted as toxic by all four models, representing the highest-confidence candidates for experimental validation. Horizontal bars (left panel) indicate the total number of positive predictions per model.

This observation reflects a broader limitation of the binary classification approach in peptide toxicity prediction. Biologically active peptides operate within a continuous, multi-label functional space: a single peptide may simultaneously exhibit antimicrobial, cytotoxic, and hemolytic activity, with varying intensities depending on context, concentration, and target cell type (Fjell and Hiss, 2012). By collapsing this diversity into a toxic/non-toxic dichotomy, datasets necessarily aggregate functionally heterogeneous subpopulations under the same negative label. Models trained under this configuration are not necessarily learning incorrectly, they are responding coherently to a real biological signal that the original annotation scheme failed to capture.

## Conclusion

We introduced NoiTox, a multi-model strategy for peptide toxicity prediction integrating three complementary architectures, evaluated as four model variants: NoiToxML (Bo/full), an Extremely Randomized Trees classifier with Boruta-selected or complete physicochemical, compositional, and atomic descriptors; NoiToxDP, a deep learning model leveraging ESM-2 protein language model embeddings; and NoiToxGP, a graph neural network that combines ESM-2 residue embeddings, a fully connected residue graph with attention, and the 44 Boruta descriptors as global features. Across the CD-HIT benchmarks and the independent dataset, the models achieved balanced discrimination between toxic and non-toxic peptides. On the CD-HIT test sets, ToxiPep retained the highest ACC and MCC, with the NoiTox consensus the closest configuration (MCC 0.666–0.680), whereas on the independent dataset the NoiToxML variants and the consensus exceeded ToxiPep in MCC (0.702–0.730 versus 0.631–0.670). Probability averaging across the four variants matched or exceeded the best individual variant, and is the strategy we recommend for end users. NoiToxML (Bo) combines interpretability, relying solely on 44 physicochemical, compositional, and atomic descriptors, with competitive performance, while the embedding-based variants provide complementary representations. This pattern underscores a central strength of the multi-model NoiTox design: by combining architectures with different inductive biases, at least one configuration approaches or exceeds state-of-the-art performance on every benchmark tested.

Feature importance analyses in NoiToxML (Bo) consistently highlighted the relevance of cysteine-related descriptors, sulfur content, dipeptide composition, and net charge, supporting established biochemical mechanisms associated with peptide reactivity and cytotoxicity. The analysis of misclassified peptides further revealed regions of overlap in feature space that help explain prediction errors and may correspond to peptides with intermediate or atypical toxicity profiles.

Twenty sequences labeled as non-toxic in the benchmark datasets were unanimously predicted as toxic by all four predictors, including ToxiPep as an external reference (Figure 6; Supplementary Table S10). Because this agreement spans architectures with different inductive biases, it is more consistent with a shared signal than with model-specific errors, and we propose these sequences as the highest-confidence candidates for experimental validation. Cross-referencing negative-labeled sequences with curated databases of hemolytic, cytotoxic, and neurotoxic peptides supports this interpretation: sequences misclassified as toxic were enriched for entries annotated as toxic in external databases (Supplementary Table S9), indicating that some apparent errors may reflect genuine biological activity in sequences whose negative label rests on the absence of annotation. However, removing the training sequences that all NoiTox models contradicted in out-of-fold predictions did not improve the ranking performance of any model and reduced the AUROC of the NoiToxML variants and of the consensus, indicating that such sequences carry useful information and should be curated rather than removed automatically. Experimental testing of the twenty candidates is therefore the natural next step. This finding underscores a fundamental challenge in computational toxicology: the negative class is typically defined by the absence of evidence rather than confirmed inactivity, introducing a persistent source of ambiguity that limits the interpretability of standard evaluation metrics. Future efforts toward multi-label annotation schemes and positive-unlabeled learning strategies may help mitigate this limitation and enable more faithful benchmarking of peptide toxicity predictors.

## Data availability

All the data and source code are available at the GitHub repository: https://github.com/goalmeida05/NoiTox

## Acknowledgments

Coordenação de Aperfeiçoamento de Pessoal de Nível Superior – Brasil (CAPES); Conselho Nacional de Desenvolvimento Científico e Tecnológico (CNPq); Fundação de Amparo à Pesquisa do Estado de Minas Gerais (FAPEMIG).

