## Supplementary Material for "NoiTox: A Machine Learning Strategy for Peptide Toxicity Prediction Reveals Label Noise in Benchmark Datasets"

### SUPPLEMENTARY MATERIALS

October 8, 2026

### 1 Model parameters and evaluation

Table S1: Hyperparameters of the NoiTox models used to generate the reported results. All models were trained with random seeds 42–46.

| Model | Parameter | Value |
|---|---|---|
| NoiToxML (Bo) / (full) | Classifier | Extremely Randomized Trees |
| | Number of trees | 500 |
| | Split criterion | Gini |
| | Features per split | $\sqrt{\text{number of features}}$ |
| | Maximum depth | unlimited |
| | Minimum samples per split / leaf | 2 / 1 |
| | Bootstrap; class weights | no; none |
| | Random state | seed |
| | Input features | 44 (Bo) / 8,461 (full) |
| NoiToxDP | Input | ESM-2 (35M, layer 12), mean-pooled, 480 dimensions |
| | Architecture | 480 $\rightarrow$ 256 (ReLU, dropout 0.3) $\rightarrow$ 1 |
| | Loss; optimizer | Binary cross-entropy; Adam |
| | Learning rate; weight decay | $10^{-3}$; 0 |
| | Batch size; maximum epochs | 64; 100 |
| | Early stopping | Validation loss, patience 10 |
| | Validation set | 10% of the training set, stratified |
| NoiToxGP | Node features | ESM-2 (35M) per-residue embeddings, 480 dimensions |
| | Architecture | 3 GATv2 layers, 128 hidden units, 4 heads, attention pooling (dimension 64), dropout 0.2 |
| | Global features | 44 Boruta descriptors $\rightarrow$ Linear–LayerNorm–ELU (64 units), concatenated with the graph representation |
| | Loss; optimizer | Binary cross-entropy; Adam |
| | Learning rate; weight decay | $10^{-3}$; 0 |
| | Batch size; maximum epochs | 32; 100 |
| | Early stopping | Validation loss, patience 15 |
| | Validation set | 10% of the training set, stratified |

Table S2: The 44 physicochemical, compositional, and atomic descriptors confirmed as informative by the Boruta feature selection algorithm, used by NoiToxML (Bo). Descriptors are grouped by category.

| Category | Descriptors |
|---|---|
| Global physicochemical (5) | Norm_mass, Norm_isoelectric_point, Norm_net_charge, Norm_gravy, Norm_hidro/total |
| Length (1) | len |
| Atomic composition (6) | normnC, normnH, normnN, normnO, normnS, total_elements |
| CKSAAGP, $k$=1 (9) | norm_hydrophobic_negatively_charged_CKSAAGP, norm_hydrophobic_polar_non_charged_CKSAAGP, norm_hydrophobic_positively_charged_CKSAAGP, norm_negatively_charged_polar_non_charged_CKSAAGP, norm_polar_non_charged_hydrophobic_CKSAAGP, norm_polar_non_charged_polar_non_charged_CKSAAGP, norm_polar_non_charged_positively_charged_CKSAAGP, norm_positively_charged_hydrophobic_CKSAAGP, norm_positively_charged_polar_non_charged_CKSAAGP |
| Amino acid composition (9) | normA, normC, normI, normK normL, normM, normT, normV, normW |
| Dipeptide composition (13) | normCC, normCG, normCK, normCP, normCS, normCY, normDC, normEC, normGC, normKK, normPC, normTC, normWC |
| Tripeptide composition (1) | normCCS |

Table S3: Comparison of NoiTox metrics with competing methods under different CD-HIT thresholds. NoiTox values are the mean ± SD over five random seeds.

| CD-HIT | Method | ACC | SN | SP | MCC |
|---|---|---|---|---|---|
| | ToxinPred2.0* | 0.596 | 0.974 | 0.217 | 0.293 |
| | ToxinPred2.0** | 0.619 | **0.975** | 0.264 | 0.340 |
| | CSM-Toxin | 0.506 | 0.062 | **0.949** | 0.025 |
| | PepNet | 0.790 | 0.677 | 0.903 | 0.596 |
| | ToxiPep | **0.850** | 0.827 | 0.873 | **0.701** |
| 0.9 | NoiToxML (full) | 0.826 ± 0.001 | 0.773 ± 0.003 | 0.879 ± 0.003 | 0.655 ± 0.003 |
| | NoiToxDP | 0.832 ± 0.004 | 0.799 ± 0.014 | 0.866 ± 0.008 | 0.666 ± 0.007 |
| | NoiToxGP | 0.827 ± 0.005 | 0.780 ± 0.019 | 0.875 ± 0.013 | 0.658 ± 0.008 |
| | NoiToxML (Bo) | 0.824 ± 0.003 | 0.772 ± 0.003 | 0.876 ± 0.003 | 0.651 ± 0.006 |
| | Consensus (prob. avg.) | 0.838 ± 0.005 | 0.788 ± 0.002 | 0.887 ± 0.010 | 0.679 ± 0.011 |
| | Consensus (majority) | 0.839 ± 0.005 | 0.788 ± 0.006 | 0.889 ± 0.007 | 0.680 ± 0.009 |
| | ToxinPred2.0* | 0.598 | 0.977 | 0.219 | 0.300 |
| | ToxinPred2.0** | 0.625 | **0.980** | 0.269 | 0.354 |
| | CSM-Toxin | 0.499 | 0.048 | **0.950** | −0.007 |
| | PepNet | 0.786 | 0.683 | 0.890 | 0.586 |
| | ToxiPep | **0.840** | 0.826 | 0.854 | **0.680** |
| 0.8 | NoiToxML (full) | 0.812 ± 0.004 | 0.763 ± 0.005 | 0.862 ± 0.005 | 0.628 ± 0.007 |
| | NoiToxDP | 0.823 ± 0.006 | 0.800 ± 0.018 | 0.845 ± 0.023 | 0.647 ± 0.012 |
| | NoiToxGP | 0.823 ± 0.005 | 0.775 ± 0.040 | 0.870 ± 0.036 | 0.649 ± 0.009 |
| | NoiToxML (Bo) | 0.820 ± 0.003 | 0.774 ± 0.004 | 0.865 ± 0.003 | 0.642 ± 0.005 |
| | Consensus (prob. avg.) | 0.833 ± 0.001 | 0.796 ± 0.009 | 0.870 ± 0.009 | 0.668 ± 0.003 |
| | Consensus (majority) | 0.832 ± 0.003 | 0.794 ± 0.011 | 0.870 ± 0.009 | 0.666 ± 0.006 |

Values in bold is the highest value for each metric at each threshold.
*ToxinPred2.0 (RF). **ToxinPred2.0 (Hybrid).
Consensus: probability averaging and majority voting over NoiToxML (full), NoiToxML (Bo), NoiToxDP and NoiToxGP.

Table S4: Comparison of NoiTox metrics with competing method on the independent dataset. NoiTox values are the mean ± SD over five random seeds.

| CD-HIT | Method | ACC | SN | SP | MCC |
|---|---|---|---|---|---|
| 0.9 | ToxiPep | 0.835 | 0.825 | 0.846 | 0.670 |
| | NoiToxML (full) | 0.849 ± 0.006 | 0.764 ± 0.009 | **0.934 ± 0.002** | 0.709 ± 0.011 |
| | NoiToxDP | 0.825 ± 0.012 | **0.846 ± 0.021** | 0.803 ± 0.006 | 0.650 ± 0.024 |
| | NoiToxGP | 0.810 ± 0.010 | 0.798 ± 0.043 | 0.822 ± 0.034 | 0.621 ± 0.020 |
| | NoiToxML (Bo) | **0.863 ± 0.003** | 0.813 ± 0.003 | 0.913 ± 0.002 | **0.730 ± 0.005** |
| | Consensus (prob. avg.) | 0.861 ± 0.005 | 0.833 ± 0.009 | 0.890 ± 0.014 | 0.724 ± 0.011 |
| | Consensus (majority) | **0.863 ± 0.008** | 0.835 ± 0.010 | 0.891 ± 0.015 | 0.727 ± 0.017 |
| 0.8 | ToxiPep | 0.815 | 0.821 | 0.810 | 0.631 |
| | NoiToxML (full) | 0.851 ± 0.003 | 0.782 ± 0.004 | **0.919 ± 0.004** | 0.708 ± 0.007 |
| | NoiToxDP | 0.818 ± 0.005 | **0.859 ± 0.019** | 0.777 ± 0.023 | 0.638 ± 0.011 |
| | NoiToxGP | 0.808 ± 0.016 | 0.793 ± 0.053 | 0.822 ± 0.060 | 0.619 ± 0.032 |
| | NoiToxML (Bo) | 0.850 ± 0.005 | 0.812 ± 0.005 | 0.887 ± 0.005 | 0.702 ± 0.009 |
| | Consensus (prob. avg.) | **0.856 ± 0.004** | 0.834 ± 0.010 | 0.878 ± 0.016 | **0.713 ± 0.009** |
| | Consensus (majority) | 0.855 ± 0.004 | 0.833 ± 0.016 | 0.878 ± 0.018 | 0.712 ± 0.009 |

Values in bold is the highest value for each metric at each threshold.
Consensus: probability averaging and majority voting over NoiToxML (full), NoiToxML (Bo), NoiToxDP and NoiToxGP.

### 2 NoiTox results compared with state-of-the-art methods

Table S5: ToxinPred2.0 (RF) performance across decision thresholds on the CD-HIT 0.8 and 0.9 test sets, reproduced locally from the standalone model. Values in bold indicate the highest ACC and MCC at each CD-HIT threshold. Decision Threshold refers to the RF probability cutoff used to classify a sequence as toxic. – indicates an undefined MCC (zero-variance confusion matrix, no true negatives predicted).

| CD-HIT Threshold | Decision Threshold | ACC | SN | SP | MCC |
|---|---|---|---|---|---|
| | 0.1 | 0.500 | 1.000 | 0.000 | – |
| | 0.2 | 0.500 | 1.000 | 0.000 | – |
| | 0.3 | 0.507 | 1.000 | 0.014 | 0.085 |
| | 0.4 | 0.538 | 0.993 | 0.082 | 0.181 |
| 0.8 | 0.5 | 0.604 | 0.976 | 0.232 | 0.310 |
| | 0.6 | 0.687 | 0.902 | 0.473 | 0.415 |
| | 0.7 | 0.736 | 0.762 | 0.709 | 0.472 |
| | 0.8 | **0.750** | 0.669 | 0.831 | **0.507** |
| | 0.9 | 0.689 | 0.463 | **0.915** | 0.423 |
| | 0.1 | 0.500 | 1.000 | 0.000 | – |
| | 0.2 | 0.501 | 1.000 | 0.002 | 0.034 |
| | 0.3 | 0.509 | 1.000 | 0.017 | 0.093 |
| | 0.4 | 0.537 | 0.996 | 0.078 | 0.185 |
| 0.9 | 0.5 | 0.601 | 0.973 | 0.229 | 0.302 |
| | 0.6 | 0.694 | 0.899 | 0.488 | 0.425 |
| | 0.7 | 0.751 | 0.767 | 0.735 | 0.502 |
| | 0.8 | **0.751** | 0.665 | 0.838 | **0.511** |
| | 0.9 | 0.690 | 0.447 | **0.932** | 0.434 |

Table S6: ToxinPred2.0 (Hybrid) performance across decision thresholds on the CD-HIT 0.8 and 0.9 test sets, reproduced locally from the standalone model. The Hybrid Score combines the ML, MERCI, and BLAST components and ranges from −0.5 to 2.0, hence the wider threshold sweep compared to the RF model (Supplementary Table S5). Bold rows indicate threshold 0.5, the value used for the main comparison in Table 2, selected for closest reproduction of the published metrics. – indicates an undefined MCC (zero-variance confusion matrix).

| CD-HIT Threshold | Decision Threshold | ACC | SN | SP | MCC |
|---|---|---|---|---|---|
| 0.8 | 0.0 | 0.504 | 1.000 | 0.007 | 0.060 |
| | 0.1 | 0.508 | 1.000 | 0.016 | 0.089 |
| | 0.2 | 0.510 | 1.000 | 0.020 | 0.101 |
| | 0.3 | 0.520 | 1.000 | 0.039 | 0.141 |
| | 0.4 | 0.548 | 0.996 | 0.101 | 0.216 |
| | **0.5** | **0.619** | **0.983** | **0.255** | **0.347** |
| | 0.6 | 0.701 | 0.914 | 0.489 | 0.444 |
| | 0.7 | 0.751 | 0.785 | 0.718 | 0.504 |
| | 0.8 | 0.767 | 0.700 | 0.834 | 0.539 |
| | 0.9 | 0.730 | 0.542 | 0.918 | 0.496 |
| | 1.0 | 0.669 | 0.353 | 0.984 | 0.435 |
| | 1.2 | 0.661 | 0.336 | 0.987 | 0.425 |
| | 1.4 | 0.632 | 0.274 | 0.990 | 0.378 |
| | 1.6 | 0.504 | 0.010 | 0.999 | 0.057 |
| | 1.8 | 0.504 | 0.010 | 0.999 | 0.057 |
| | 2.0 | 0.500 | 0.000 | 1.000 | – |
| 0.9 | 0.0 | 0.503 | 1.000 | 0.006 | 0.053 |
| | 0.1 | 0.505 | 1.000 | 0.009 | 0.068 |
| | 0.2 | 0.507 | 1.000 | 0.015 | 0.086 |
| | 0.3 | 0.516 | 1.000 | 0.033 | 0.129 |
| | 0.4 | 0.549 | 0.998 | 0.101 | 0.223 |
| | **0.5** | **0.617** | **0.975** | **0.259** | **0.336** |
| | 0.6 | 0.711 | 0.908 | 0.513 | 0.459 |
| | 0.7 | 0.767 | 0.783 | 0.752 | 0.535 |
| | 0.8 | 0.771 | 0.692 | 0.849 | 0.548 |
| | 0.9 | 0.732 | 0.532 | 0.931 | 0.505 |
| | 1.0 | 0.671 | 0.357 | 0.985 | 0.440 |
| | 1.2 | 0.668 | 0.342 | 0.993 | 0.442 |
| | 1.4 | 0.633 | 0.270 | 0.997 | 0.388 |
| | 1.6 | 0.502 | 0.003 | 1.000 | 0.041 |
| | 1.8 | 0.502 | 0.003 | 1.000 | 0.041 |
| | 2.0 | 0.500 | 0.000 | 1.000 | – |

Table S7: Threshold-independent discrimination performance (AUROC and AUPRC) for all models with sequence-level scores available, on the CD-HIT 0.8/0.9 test sets and on the independent dataset (evaluated separately for the two CD-HIT-trained NoiTox model groups). NoiTox values are mean ± SD over five random seeds; the consensus row refers to probability averaging (majority voting has no continuous score). CSM-Toxin is excluded because its reported score reflects confidence in the predicted class rather than a consistent probability of toxicity.

| Benchmark | Method | AUROC | AUPRC |
|---|---|---|---|
| Test 0.8 | ToxiPep | 0.907 | 0.919 |
| | PepNet | 0.829 | 0.865 |
| | ToxinPred2.0 (RF) | 0.812 | 0.793 |
| | ToxinPred2.0 (Hybrid) | 0.844 | 0.842 |
| | NoiToxML (full) | 0.892 ± 0.001 | 0.904 ± 0.001 |
| | NoiToxML (Bo) | 0.895 ± 0.000 | 0.908 ± 0.000 |
| | NoiToxDP | 0.903 ± 0.003 | 0.914 ± 0.002 |
| | NoiToxGP | 0.903 ± 0.003 | 0.905 ± 0.002 |
| | Consensus (prob. avg.) | **0.913 ± 0.001** | **0.923 ± 0.001** |
| Test 0.9 | ToxiPep | 0.919 | 0.926 |
| | PepNet | 0.843 | 0.875 |
| | ToxinPred2.0 (RF) | 0.825 | 0.818 |
| | ToxinPred2.0 (Hybrid) | 0.853 | 0.861 |
| | NoiToxML (full) | 0.905 ± 0.001 | 0.908 ± 0.001 |
| | NoiToxML (Bo) | 0.904 ± 0.001 | 0.912 ± 0.001 |
| | NoiToxDP | 0.921 ± 0.001 | 0.928 ± 0.001 |
| | NoiToxGP | 0.911 ± 0.006 | 0.916 ± 0.009 |
| | Consensus (prob. avg.) | **0.926 ± 0.001** | **0.932 ± 0.002** |
| Independent (0.8) | ToxiPep | 0.889 | 0.906 |
| | NoiToxML (full) | 0.926 ± 0.001 | 0.931 ± 0.001 |
| | NoiToxML (Bo) | 0.912 ± 0.001 | 0.907 ± 0.001 |
| | NoiToxDP | 0.903 ± 0.009 | 0.913 ± 0.010 |
| | NoiToxGP | 0.889 ± 0.014 | 0.900 ± 0.007 |
| | Consensus (prob. avg.) | **0.931 ± 0.003** | **0.940 ± 0.002** |
| Independent (0.9) | ToxiPep | 0.886 | 0.883 |
| | NoiToxML (full) | **0.934 ± 0.001** | 0.934 ± 0.002 |
| | NoiToxML (Bo) | 0.918 ± 0.001 | 0.906 ± 0.001 |
| | NoiToxDP | 0.904 ± 0.006 | 0.915 ± 0.007 |
| | NoiToxGP | 0.882 ± 0.013 | 0.887 ± 0.015 |
| | Consensus (prob. avg.) | 0.932 ± 0.003 | **0.935 ± 0.004** |

Table S8: Pairwise comparisons between models with sequence-level predictions on each benchmark. $\Delta$MCC = MCC$_A$ − MCC$_B$, reported as mean ± SD over the five random seeds of the NoiTox models (ToxiPep and PepNet are single, deterministic evaluations). 95% CI: mean over seeds of the bootstrap bounds (10,000 resamples). McNemar $p$: median two-sided $p$ over seeds. Holm $p$: Holm-adjusted value of the median $p$ across all pairs within each benchmark (bold: $< 0.05$). Seeds: number of seeds (of 5) in which the McNemar test ($p < 0.05$) / the bootstrap CI (excluding zero) indicated a significant difference. The consensus is the probability-averaging strategy. PepNet was not evaluated on the independent dataset.

| Comparison (A vs B) | ΔMCC | 95% CI | McNemar $p$ | Holm $p$ | Seeds |
|---|---|---|---|---|---|
| *Test set, CD-HIT 0.8 ($n$ = 1,388)* | | | | | |
| ToxiPep vs PepNet | +0.094 ± 0.000 | [+0.048, +0.141] | <0.001 | **<0.001** | 5/5 |
| ToxiPep vs NoiToxML (full) | +0.052 ± 0.007 | [+0.017, +0.088] | 0.002 | **0.046** | 5/5 |
| ToxiPep vs NoiToxML (Bo) | +0.038 ± 0.005 | [+0.003, +0.073] | 0.022 | 0.307 | 3/3 |
| ToxiPep vs NoiToxDP | +0.034 ± 0.012 | [−0.001, +0.068] | 0.059 | 0.646 | 2/1 |
| ToxiPep vs NoiToxGP | +0.031 ± 0.009 | [−0.006, +0.068] | 0.037 | 0.446 | 3/2 |
| ToxiPep vs Consensus (prob. avg.) | +0.012 ± 0.003 | [−0.020, +0.044] | 0.433 | 1.000 | 0/0 |
| PepNet vs NoiToxML (full) | −0.042 ± 0.007 | [−0.087, +0.001] | 0.030 | 0.396 | 4/2 |
| PepNet vs NoiToxML (Bo) | −0.056 ± 0.005 | [−0.101, −0.010] | 0.007 | 0.108 | 5/5 |
| PepNet vs NoiToxDP | −0.060 ± 0.012 | [−0.109, −0.012] | 0.004 | 0.070 | 5/4 |
| PepNet vs NoiToxGP | −0.063 ± 0.009 | [−0.109, −0.017] | 0.005 | 0.082 | 5/5 |
| PepNet vs Consensus (prob. avg.) | −0.082 ± 0.003 | [−0.127, −0.037] | <0.001 | **0.002** | 5/5 |
| NoiToxML (full) vs NoiToxML (Bo) | −0.014 ± 0.009 | [−0.042, +0.012] | 0.289 | 1.000 | 0/0 |
| NoiToxML (full) vs NoiToxDP | −0.018 ± 0.011 | [−0.053, +0.016] | 0.178 | 1.000 | 0/0 |
| NoiToxML (full) vs NoiToxGP | −0.021 ± 0.006 | [−0.057, +0.014] | 0.304 | 1.000 | 0/0 |
| NoiToxML (full) vs Consensus (prob. avg.) | −0.040 ± 0.006 | [−0.068, −0.014] | 0.004 | 0.070 | 5/5 |
| NoiToxML (Bo) vs NoiToxDP | −0.004 ± 0.014 | [−0.040, +0.031] | 0.468 | 1.000 | 0/0 |
| NoiToxML (Bo) vs NoiToxGP | −0.007 ± 0.008 | [−0.044, +0.030] | 0.644 | 1.000 | 0/0 |
| NoiToxML (Bo) vs Consensus (prob. avg.) | −0.026 ± 0.006 | [−0.054, +0.003] | 0.086 | 0.859 | 1/2 |
| NoiToxDP vs NoiToxGP | −0.003 ± 0.012 | [−0.033, +0.028] | 0.839 | 1.000 | 0/0 |
| NoiToxDP vs Consensus (prob. avg.) | −0.022 ± 0.011 | [−0.046, +0.002] | 0.115 | 0.919 | 1/1 |
| NoiToxGP vs Consensus (prob. avg.) | −0.019 ± 0.007 | [−0.045, +0.007] | 0.092 | 0.859 | 1/0 |
| *Test set, CD-HIT 0.9 ($n$ = 1,766)* | | | | | |
| ToxiPep vs PepNet | +0.104 ± 0.000 | [+0.064, +0.145] | <0.001 | **<0.001** | 5/5 |
| ToxiPep vs NoiToxML (full) | +0.046 ± 0.003 | [+0.018, +0.074] | 0.001 | **0.019** | 5/5 |
| ToxiPep vs NoiToxML (Bo) | +0.049 ± 0.006 | [+0.020, +0.080] | <0.001 | **0.009** | 5/5 |
| ToxiPep vs NoiToxDP | +0.034 ± 0.007 | [+0.002, +0.067] | 0.049 | 0.494 | 3/3 |
| ToxiPep vs NoiToxGP | +0.043 ± 0.008 | [+0.011, +0.076] | 0.007 | 0.092 | 5/5 |
| ToxiPep vs Consensus (prob. avg.) | +0.022 ± 0.011 | [−0.007, +0.051] | 0.169 | 1.000 | 1/1 |
| PepNet vs NoiToxML (full) | −0.059 ± 0.003 | [−0.096, −0.021] | <0.001 | **0.007** | 5/5 |
| PepNet vs NoiToxML (Bo) | −0.055 ± 0.006 | [−0.095, −0.014] | 0.003 | **0.036** | 5/5 |
| PepNet vs NoiToxDP | −0.070 ± 0.007 | [−0.111, −0.029] | <0.001 | **0.002** | 5/5 |
| PepNet vs NoiToxGP | −0.061 ± 0.008 | [−0.102, −0.020] | 0.001 | **0.019** | 5/5 |
| PepNet vs Consensus (prob. avg.) | −0.083 ± 0.011 | [−0.122, −0.043] | <0.001 | **<0.001** | 5/5 |
| NoiToxML (full) vs NoiToxML (Bo) | +0.004 ± 0.006 | [−0.018, +0.026] | 0.614 | 1.000 | 0/0 |
| NoiToxML (full) vs NoiToxDP | −0.011 ± 0.008 | [−0.042, +0.019] | 0.378 | 1.000 | 0/0 |
| NoiToxML (full) vs NoiToxGP | −0.002 ± 0.009 | [−0.032, +0.027] | 0.659 | 1.000 | 0/0 |
| NoiToxML (full) vs Consensus (prob. avg.) | −0.024 ± 0.013 | [−0.047, −0.001] | 0.019 | 0.227 | 3/3 |
| NoiToxML (Bo) vs NoiToxDP | −0.015 ± 0.011 | [−0.045, +0.015] | 0.404 | 1.000 | 1/0 |
| NoiToxML (Bo) vs NoiToxGP | −0.006 ± 0.014 | [−0.036, +0.023] | 0.563 | 1.000 | 0/0 |
| NoiToxML (Bo) vs Consensus (prob. avg.) | −0.028 ± 0.011 | [−0.050, −0.006] | 0.028 | 0.309 | 4/4 |

| Comparison (A vs B) | ΔMCC | 95% CI | McNemar $p$ | Holm $p$ | Seeds |
|---|---|---|---|---|---|
| NoiToxDP vs NoiToxGP | $+0.009 \pm 0.006$ | $[-0.019, +0.037]$ | 0.594 | 1.000 | 0/0 |
| NoiToxDP vs Consensus (prob. avg.) | $-0.013 \pm 0.009$ | $[-0.036, +0.011]$ | 0.481 | 1.000 | 0/0 |
| NoiToxGP vs Consensus (prob. avg.) | $-0.021 \pm 0.014$ | $[-0.043, -0.000]$ | 0.083 | 0.744 | 2/3 |
| *Independent dataset, models trained at CD-HIT 0.8 ($n = 958$)* | | | | | |
| ToxiPep vs NoiToxML (full) | $-0.077 \pm 0.007$ | $[-0.126, -0.029]$ | 0.006 | 0.072 | 5/5 |
| ToxiPep vs NoiToxML (Bo) | $-0.071 \pm 0.009$ | $[-0.120, -0.023]$ | 0.006 | 0.072 | 5/5 |
| ToxiPep vs NoiToxDP | $-0.008 \pm 0.011$ | $[-0.056, +0.040]$ | 0.806 | 1.000 | 0/0 |
| ToxiPep vs NoiToxGP | $+0.012 \pm 0.032$ | $[-0.040, +0.063]$ | 0.618 | 1.000 | 1/1 |
| ToxiPep vs Consensus (prob. avg.) | $-0.083 \pm 0.009$ | $[-0.128, -0.037]$ | <0.001 | **0.004** | 5/5 |
| NoiToxML (full) vs NoiToxML (Bo) | $+0.006 \pm 0.016$ | $[-0.031, +0.043]$ | 0.590 | 1.000 | 0/0 |
| NoiToxML (full) vs NoiToxDP | $+0.069 \pm 0.015$ | $[+0.018, +0.120]$ | 0.013 | 0.114 | 4/4 |
| NoiToxML (full) vs NoiToxGP | $+0.089 \pm 0.033$ | $[+0.038, +0.141]$ | 0.013 | 0.114 | 5/5 |
| NoiToxML (full) vs Consensus (prob. avg.) | $-0.006 \pm 0.012$ | $[-0.043, +0.032]$ | 0.828 | 1.000 | 0/0 |
| NoiToxML (Bo) vs NoiToxDP | $+0.063 \pm 0.009$ | $[+0.012, +0.116]$ | 0.019 | 0.135 | 4/4 |
| NoiToxML (Bo) vs NoiToxGP | $+0.083 \pm 0.038$ | $[+0.032, +0.134]$ | 0.006 | 0.072 | 4/4 |
| NoiToxML (Bo) vs Consensus (prob. avg.) | $-0.011 \pm 0.014$ | $[-0.050, +0.027]$ | 0.640 | 1.000 | 0/0 |
| NoiToxDP vs NoiToxGP | $+0.020 \pm 0.038$ | $[-0.025, +0.064]$ | 0.379 | 1.000 | 1/1 |
| NoiToxDP vs Consensus (prob. avg.) | $-0.075 \pm 0.012$ | $[-0.114, -0.036]$ | <0.001 | **0.004** | 5/5 |
| NoiToxGP vs Consensus (prob. avg.) | $-0.095 \pm 0.029$ | $[-0.134, -0.056]$ | <0.001 | **<0.001** | 5/5 |
| *Independent dataset, models trained at CD-HIT 0.9 ($n = 958$)* | | | | | |
| ToxiPep vs NoiToxML (full) | $-0.038 \pm 0.011$ | $[-0.085, +0.007]$ | 0.305 | 0.916 | 0/2 |
| ToxiPep vs NoiToxML (Bo) | $-0.060 \pm 0.005$ | $[-0.107, -0.014]$ | 0.021 | 0.206 | 5/5 |
| ToxiPep vs NoiToxDP | $+0.020 \pm 0.024$ | $[-0.030, +0.071]$ | 0.379 | 0.916 | 0/1 |
| ToxiPep vs NoiToxGP | $+0.049 \pm 0.020$ | $[-0.004, +0.101]$ | 0.138 | 0.827 | 2/1 |
| ToxiPep vs Consensus (prob. avg.) | $-0.054 \pm 0.011$ | $[-0.099, -0.009]$ | 0.029 | 0.262 | 3/3 |
| NoiToxML (full) vs NoiToxML (Bo) | $-0.021 \pm 0.007$ | $[-0.057, +0.015]$ | 0.161 | 0.827 | 0/0 |
| NoiToxML (full) vs NoiToxDP | $+0.059 \pm 0.023$ | $[+0.010, +0.109]$ | 0.077 | 0.587 | 2/3 |
| NoiToxML (full) vs NoiToxGP | $+0.088 \pm 0.030$ | $[+0.038, +0.138]$ | 0.013 | 0.146 | 4/5 |
| NoiToxML (full) vs Consensus (prob. avg.) | $-0.015 \pm 0.015$ | $[-0.051, +0.021]$ | 0.207 | 0.828 | 1/0 |
| NoiToxML (Bo) vs NoiToxDP | $+0.080 \pm 0.022$ | $[+0.030, +0.130]$ | 0.006 | 0.076 | 4/5 |
| NoiToxML (Bo) vs NoiToxGP | $+0.109 \pm 0.025$ | $[+0.058, +0.160]$ | <0.001 | **0.004** | 5/5 |
| NoiToxML (Bo) vs Consensus (prob. avg.) | $+0.006 \pm 0.012$ | $[-0.030, +0.043]$ | 0.826 | 0.916 | 0/0 |
| NoiToxDP vs NoiToxGP | $+0.029 \pm 0.032$ | $[-0.015, +0.073]$ | 0.073 | 0.587 | 2/2 |
| NoiToxDP vs Consensus (prob. avg.) | $-0.074 \pm 0.016$ | $[-0.113, -0.035]$ | <0.001 | **0.008** | 5/5 |
| NoiToxGP vs Consensus (prob. avg.) | $-0.103 \pm 0.019$ | $[-0.142, -0.064]$ | <0.001 | **<0.001** | 5/5 |

Table S9: Fisher's exact test (one-sided) for enrichment of toxicity-database matches among misclassified negative-labeled sequences (false positives) relative to correctly classified negative-labeled sequences (true negatives), by model and CD-HIT threshold. Rate: proportion of sequences in that group matching at least one of the DBAASP, Hemolytic2, UniProt (cytotoxic), or UniProt (neurotoxic) databases. OR: odds ratio. Bold: $p < 0.05$.

| CD-HIT | Method | Misclassified (matched/n, rate) | Correct (matched/n, rate) | OR | $p$ |
|---|---|---|---|---|---|
| 0.8 | NoiToxML (full) | 1/22 (4.55%) | 9/672 (1.34%) | 3.51 | 0.277 |
| | NoiToxML (Bo) | 1/34 (2.94%) | 9/660 (1.36%) | 2.19 | 0.397 |
| | NoiToxDP | 9/121 (7.44%) | 1/573 (0.17%) | 45.96 | **<0.001** |
| | NoiToxGP | 10/101 (9.90%) | 0/593 (0.00%) | ∞ | **<0.001** |
| 0.9 | NoiToxML (full) | 5/96 (5.21%) | 3/787 (0.38%) | 14.36 | **0.001** |
| | NoiToxML (Bo) | 3/77 (3.90%) | 5/806 (0.62%) | 6.49 | **0.026** |
| | NoiToxDP | 7/118 (5.93%) | 1/765 (0.13%) | 48.18 | **<0.001** |
| | NoiToxGP | 8/185 (4.32%) | 0/698 (0.00%) | ∞ | **<0.001** |

Table S10: The 20 sequences labeled non-toxic in the benchmark datasets that were unanimously predicted as toxic by all four evaluated models (NoiToxML (Bo), NoiToxDP, NoiToxGP, and ToxiPep as an external reference), corresponding to the dark red bar ($n = 20$). These represent the highest-confidence candidates for experimental validation discussed in Section Incorrect predictions.

| # | Sequence | ID |
|---|---|---|
| 1 | TDCGGKTCSEAQVCKDGKCVCVIGQCRKYCPNGFKKDENGCTFPCTCA | P81499 |
| 2 | YSRCQLQGFNCVVRSYGLPTIPCCRGLTCRSYFPGSTYGRCQRF | P0C1Z7 |
| 3 | QTCASRCPRPCNAGLCCSIYGYCGSGNAYCGAGNCRCQCRG | P83596 |
| 4 | DICDIAIAQCSLTLCQDCENTPICELAVKGSCPPPWS | P58548 |
| 5 | GLFLNTVKDVAKDVAKDVAGKLLESLKCKITGCKP | P0DQK4 |
| 6 | GILLDKLKNFAKTAGKGVLQSLLNTASCKLSGQC | P40839 |
| 7 | GIFLNALKNFAKTAGKGVLQSVLNTASCKLSKQC | C0HL00 |
| 8 | GLWKTLKDSAKSAVTNVAVTMLDKLRCKLTGGC | C0HL04 |
| 9 | GLPTCGETCFGGTCNTPGCTCDSSWPICTHN | P85186 |
| 10 | GSIPACGESCFKGKCYTPGCTCSKYPLCAKN | C0HKK2 |
| 11 | GLPTCGETCFGGTCNTPGCSCSSWPICTRD | P85128 |
| 12 | GFPTCGETCTLGTCNTPGCTCSWPICTRN | C0HKJ7 |
| 13 | GLPVCGESCFGGSCYTPGCSCTWPICTRD | P85133 |
| 14 | GDLPVCGETCFGGTCNTPGCVCAWPVCTR | C0HL29 |
| 15 | GKPICGETCAKGKCYTPKCTCNWPICYKN | C0HKJ8 |
| 16 | GLPVCGETCFGGTCNTPGCSCTDPICTRD | P85129 |
| 17 | GLPICGETCFTGTCNTPGCSCTYPICTRD | C0HL24 |
| 18 | RRRRRRRRHRRRRGRRGRRSRGRR | P83215 |
| 19 | FLPAIVGAAAKFLPKIFCVISKKC | P82837 |
| 20 | AIMDTIKDTAKTVAVGLLNKLKCKITGC | P85058 |

Table S11: Performance of the NoiTox models before and after retraining, on the CD-HIT 0.8/0.9 test sets (1,388 and 1,766 peptides) and on the independent dataset (958 peptides), all balanced between toxic and non-toxic; for the independent dataset, the threshold refers to the training partition. *Before*: previous version of the models. *After*: retrained models (Supplementary Table S1); mean ± SD over five random seeds. ACC, accuracy; SN, sensitivity; SP, specificity; MCC, Matthews correlation coefficient; decision threshold 0.5.

| CD-HIT threshold | Method | Version | ACC | SN | SP | MCC |
|---|---|---|---|---|---|---|
| *Test sets* | | | | | | |
| 0.8 | NoiToxML (full) | Before | 0.830 | 0.838 | 0.822 | 0.641 |
| | | After | 0.812 ± 0.004 | 0.763 ± 0.005 | 0.862 ± 0.005 | 0.628 ± 0.007 |
| | NoiToxML (Bo) | Before | 0.818 | 0.775 | 0.860 | 0.639 |
| | | After | 0.820 ± 0.003 | 0.774 ± 0.004 | 0.865 ± 0.003 | 0.642 ± 0.005 |
| | NoiToxDP | Before | 0.824 | 0.812 | 0.837 | 0.649 |
| | | After | 0.823 ± 0.006 | 0.800 ± 0.018 | 0.845 ± 0.023 | 0.647 ± 0.012 |
| | NoiToxGP | Before | 0.837 | 0.806 | 0.869 | 0.676 |
| | | After | 0.823 ± 0.005 | 0.775 ± 0.040 | 0.870 ± 0.036 | 0.649 ± 0.009 |
| 0.9 | NoiToxML (full) | Before | 0.818 | 0.762 | 0.873 | 0.639 |
| | | After | 0.826 ± 0.001 | 0.773 ± 0.003 | 0.879 ± 0.003 | 0.655 ± 0.003 |
| | NoiToxML (Bo) | Before | 0.823 | 0.772 | 0.876 | 0.651 |
| | | After | 0.824 ± 0.003 | 0.772 ± 0.003 | 0.876 ± 0.003 | 0.651 ± 0.006 |
| | NoiToxDP | Before | 0.838 | 0.804 | 0.874 | 0.679 |
| | | After | 0.832 ± 0.004 | 0.799 ± 0.014 | 0.866 ± 0.008 | 0.666 ± 0.007 |
| | NoiToxGP | Before | 0.830 | 0.861 | 0.798 | 0.661 |
| | | After | 0.827 ± 0.005 | 0.780 ± 0.019 | 0.875 ± 0.013 | 0.658 ± 0.008 |
| *Independent dataset* | | | | | | |
| 0.8 | NoiToxML (full) | Before | 0.811 | 0.813 | 0.809 | 0.629 |
| | | After | 0.851 ± 0.003 | 0.782 ± 0.004 | 0.919 ± 0.004 | 0.708 ± 0.007 |
| | NoiToxML (Bo) | Before | 0.847 | 0.830 | 0.864 | 0.694 |
| | | After | 0.850 ± 0.005 | 0.812 ± 0.005 | 0.887 ± 0.005 | 0.702 ± 0.009 |
| | NoiToxDP | Before | 0.803 | 0.873 | 0.729 | 0.610 |
| | | After | 0.818 ± 0.005 | 0.859 ± 0.019 | 0.777 ± 0.023 | 0.638 ± 0.011 |
| | NoiToxGP | Before | 0.816 | 0.857 | 0.774 | 0.634 |
| | | After | 0.808 ± 0.016 | 0.793 ± 0.053 | 0.822 ± 0.060 | 0.619 ± 0.032 |
| 0.9 | NoiToxML (full) | Before | 0.800 | 0.788 | 0.812 | 0.609 |
| | | After | 0.849 ± 0.006 | 0.764 ± 0.009 | 0.934 ± 0.002 | 0.709 ± 0.011 |
| | NoiToxML (Bo) | Before | 0.851 | 0.832 | 0.870 | 0.703 |
| | | After | 0.863 ± 0.003 | 0.813 ± 0.003 | 0.913 ± 0.002 | 0.730 ± 0.005 |
| | NoiToxDP | Before | 0.843 | 0.875 | 0.806 | 0.683 |
| | | After | 0.825 ± 0.012 | 0.846 ± 0.021 | 0.803 ± 0.006 | 0.650 ± 0.024 |
| | NoiToxGP | Before | 0.810 | 0.924 | 0.691 | 0.635 |
| | | After | 0.810 ± 0.010 | 0.798 ± 0.043 | 0.822 ± 0.034 | 0.621 ± 0.020 |

#### 3 Consensus strategies and recommended use

NoiTox combines the four models (NoiToxML-Full, NoiToxML-Boruta, NoiToxDP and NoiToxGP) into two consensus strategies, computed for every sequence predicted by all four models. In *probability averaging*, the raw scores of the four models are averaged and the sequence is classified as toxic if the mean is at least 0.5. In *majority voting*, each model casts a binary vote at the 0.5 threshold, the number of toxic votes is recorded (from 0 to 4), and a sequence is classified as toxic when at least three models vote toxic; the 2–2 ties that can occur with an even number of models are resolved by the probability-averaged prediction. Two flags mark unanimous calls (`unanimous_toxic`, four toxic votes; `unanimous_nontoxic`, zero toxic votes), and, when reference labels are supplied, the output also reports them (`real_label`) together with the individual score and prediction of each model. The two strategies performed almost identically (Table S12). We recommend probability averaging as the default for end users: with four models it has no ties and yields a continuous score that can be ranked and thresholded, whereas majority voting depends on a tie-breaking rule. The vote count and the unanimity flags are useful as a measure of agreement between models with different inductive biases: unanimous predictions indicate full agreement, whereas 2–2 splits identify sequences on which the models disagree and that merit closer inspection. The scores are averaged as produced by each model, without recalibration, so the mean should be interpreted as an ensemble score rather than as a calibrated probability. Sequences that all models unanimously assign to the class opposite to a reference label are the candidates for curation discussed in the main-text section on persistently misclassified sequences.

#### 4 Consensus-based cleaning of the training data

Because the consensus among NoiTox models flags sequences whose labels the models systematically contradict, we asked whether removing them from the training sets improves predictive performance. Candidate label-noise sequences were identified from out-of-fold predictions (10 stratified folds shared across models, so that every sequence was predicted only by models that had not been trained on it) as those that all four predictors (NoiToxML-Full, NoiToxML-Boruta, NoiToxDP and NoiToxGP) assigned to the class opposite to the original label. This flagged 543 of 5,544 sequences at CD-HIT 0.8 (9.8%; 349 labelled toxic and 194 non-toxic) and 544 of 7,056 at CD-HIT 0.9 (7.7%; 344 labelled toxic and 200 non-toxic). All models (original, cleaned and random-removal conditions) were then retrained under the same training configuration used for the models reported in the main text, and evaluated on the unmodified test sets and on the independent set. The original condition corresponds to a single training run (seed 42), whereas main-text values are means over five seeds. No test sequence was removed, since filtering test sequences by model errors would inflate performance by construction. To control for the effect of training on

Table S12: Performance of consensus strategies (probability averaging and majority voting across NoiToxML (full), NoiToxML (Bo), NoiToxDP, and NoiToxGP) compared to the best individual NoiTox variant, on the CD-HIT 0.8/0.9 test sets and on the independent dataset. Bold: best value per metric within each benchmark. NoiTox values come from a single training run (seed 42).

| Benchmark | Strategy | ACC | SN | SP | MCC |
|---|---|---|---|---|---|
| Test 0.8 | Best individual (NoiToxML Bo) | 0.823 | 0.780 | **0.866** | 0.648 |
| | Probability averaging | 0.834 | **0.829** | 0.840 | 0.669 |
| | Majority voting | **0.835** | **0.829** | 0.841 | **0.670** |
| Test 0.9 | Best individual (NoiToxDP) | 0.829 | **0.796** | 0.862 | 0.659 |
| | Probability averaging | **0.842** | **0.796** | **0.888** | **0.687** |
| | Majority voting | 0.841 | 0.795 | **0.888** | 0.686 |
| Independent (0.8) | Best individual (NoiToxML full) | **0.855** | 0.785 | **0.925** | **0.717** |
| | Probability averaging | 0.849 | **0.894** | 0.804 | 0.700 |
| | Majority voting | 0.851 | **0.894** | 0.808 | 0.704 |
| Independent (0.9) | Best individual (NoiToxML Bo) | 0.860 | 0.810 | **0.910** | 0.724 |
| | Probability averaging | **0.863** | **0.831** | 0.896 | **0.728** |
| | Majority voting | **0.863** | **0.831** | 0.896 | **0.728** |

less data, we repeated the procedure ten times removing, instead, the same number of randomly chosen non-flagged sequences per class, matched on length distribution.

Cleaning did not improve the ranking performance (AUROC and AUPRC) of any model (Table S13; Tables S14 and S15). Relative to random removal, it reduced the AUROC of NoiToxML (Full and Boruta) and of the probability-averaging consensus in all four evaluations ($\Delta$AUROC from $-0.004$ to $-0.027$), and the AUPRC by up to 0.066 on the independent sets. On these sets, the MCC of the same models, and of the majority-vote consensus, decreased by 0.020–0.054, reflecting a loss of specificity ($-0.046$ to $-0.076$) only partly offset by a gain in sensitivity ($+0.006$ to $+0.044$). In contrast, the AUROC of NoiToxDP and NoiToxGP was essentially unchanged ($|\Delta\text{AUROC}| \leq 0.008$), and the modest MCC gains observed for these models ($+0.005$ to $+0.035$) were not accompanied by better ranking performance, indicating a shift in operating point at the 0.5 threshold rather than improved discrimination. Random removal left the AUROC of the ML models practically unchanged relative to the original training sets ($|\Delta| \leq 0.003$), so the loss can be attributed to the flagged sequences themselves rather than to the smaller training set.

These results indicate that sequences on which all NoiTox models contradict the label are not equivalent to randomly distributed noise. A plausible reading, which we did not test directly, is that many of them are informative examples (hard or atypical peptides near the decision boundary, or under-represented regions of sequence space) whose removal reduces the coverage of the training data; the effects on MCC, AUPRC and specificity were larger on the independent sets than on the CD-HIT test sets. The greater susceptibility of the ML models,

Table S13: Change in performance after removing consensus-flagged sequences from the training set, relative to the mean of ten random-removal controls (cleaned − random). "Test" denotes the test set of the corresponding CD-HIT threshold; "Ind." denotes the independent set, evaluated with models trained at that threshold. AUROC and AUPRC are not defined for the majority vote.

| Model | Test 0.8 | Ind. 0.8 | Test 0.9 | Ind. 0.9 |
|---|---|---|---|---|
| *(A)* $\Delta MCC$ | | | | |
| NoiToxML (Full) | −0.020 | −0.043 | −0.013 | −0.043 |
| NoiToxML (Boruta) | −0.006 | −0.020 | −0.002 | −0.054 |
| NoiToxDP | +0.009 | +0.007 | +0.013 | +0.005 |
| NoiToxGP | +0.015 | +0.026 | +0.011 | +0.035 |
| Consensus (prob. avg.) | −0.003 | −0.038 | −0.001 | −0.022 |
| Consensus (majority vote) | −0.005 | −0.032 | −0.003 | −0.024 |
| *(B)* $\Delta AUROC$ | | | | |
| NoiToxML (Full) | −0.018 | −0.025 | −0.016 | −0.027 |
| NoiToxML (Boruta) | −0.004 | −0.007 | −0.007 | −0.017 |
| NoiToxDP | −0.002 | +0.007 | −0.008 | +0.002 |
| NoiToxGP | −0.004 | −0.008 | −0.002 | +0.001 |
| Consensus (prob. avg.) | −0.016 | −0.021 | −0.018 | −0.019 |
| *(C)* $\Delta AUPRC$ | | | | |
| NoiToxML (Full) | −0.014 | −0.059 | −0.017 | −0.066 |
| NoiToxML (Boruta) | −0.007 | −0.026 | −0.008 | −0.041 |
| NoiToxDP | +0.000 | +0.002 | −0.007 | −0.008 |
| NoiToxGP | +0.001 | −0.050 | −0.002 | −0.015 |
| Consensus (prob. avg.) | −0.013 | −0.054 | −0.015 | −0.050 |

which rely on hand-crafted descriptors learned only from the training data, compared with NoiToxDP and NoiToxGP, which build on pre-trained ESM-2 representations, may likewise reflect a lower dependence of the latter on the local density of training examples. We therefore recommend using the consensus as a list of candidates for expert curation (for example, literature review or experimental validation) rather than as an automatic filter.

Table S14: Change in sensitivity (SN) and specificity (SP) at the 0.5 threshold after removing consensus-flagged sequences, relative to the mean of ten random-removal controls (cleaned − random; SN / SP).

| Model | Test 0.8 | Ind. 0.8 | Test 0.9 | Ind. 0.9 |
|---|---|---|---|---|
| NoiToxML (Full) | +0.011 / −0.028 | +0.044 / −0.076 | +0.005 / −0.017 | +0.022 / −0.055 |
| NoiToxML (Boruta) | +0.012 / −0.016 | +0.033 / −0.050 | +0.006 / −0.007 | +0.006 / −0.056 |
| NoiToxDP | +0.020 / −0.009 | +0.004 / +0.003 | +0.017 / −0.003 | +0.014 / −0.009 |
| NoiToxGP | +0.053 / −0.033 | +0.084 / −0.060 | +0.027 / −0.013 | +0.031 / +0.005 |
| Consensus (prob. avg.) | +0.026 / −0.026 | +0.038 / −0.076 | +0.019 / −0.017 | +0.027 / −0.046 |
| Consensus (majority vote) | +0.023 / −0.027 | +0.042 / −0.074 | +0.016 / −0.017 | +0.024 / −0.047 |

Table S15: MCC and AUROC of models trained on the original training set (Orig.), on the training set without consensus-flagged sequences (Clean), and on training sets from which the same number of randomly chosen sequences was removed (Random; mean ± SD over ten repetitions). All evaluations use unmodified test sets. AUROC is not defined for the majority vote.

| | MCC | | | AUROC | | |
|---|---|---|---|---|---|---|
| Model | Orig. | Clean | Random | Orig. | Clean | Random |
| *Test set, CD-HIT 0.8* | | | | | | |
| NoiToxML (Full) | 0.626 | 0.611 | 0.631 ± 0.007 | 0.892 | 0.874 | 0.892 ± 0.001 |
| NoiToxML (Boruta) | 0.648 | 0.634 | 0.640 ± 0.009 | 0.895 | 0.889 | 0.893 ± 0.001 |
| NoiToxDP | 0.628 | 0.654 | 0.645 ± 0.005 | 0.904 | 0.900 | 0.902 ± 0.003 |
| NoiToxGP | 0.610 | 0.660 | 0.645 ± 0.010 | 0.903 | 0.896 | 0.900 ± 0.004 |
| Consensus (prob. avg.) | 0.669 | 0.660 | 0.663 ± 0.009 | 0.913 | 0.897 | 0.912 ± 0.001 |
| Consensus (majority vote) | 0.670 | 0.659 | 0.665 ± 0.007 | – | – | – |
| *Independent set, models trained at CD-HIT 0.8* | | | | | | |
| NoiToxML (Full) | 0.717 | 0.647 | 0.690 ± 0.009 | 0.927 | 0.899 | 0.924 ± 0.002 |
| NoiToxML (Boruta) | 0.691 | 0.674 | 0.695 ± 0.006 | 0.911 | 0.905 | 0.911 ± 0.002 |
| NoiToxDP | 0.641 | 0.651 | 0.644 ± 0.026 | 0.912 | 0.909 | 0.903 ± 0.010 |
| NoiToxGP | 0.564 | 0.628 | 0.602 ± 0.041 | 0.904 | 0.870 | 0.878 ± 0.016 |
| Consensus (prob. avg.) | 0.700 | 0.682 | 0.720 ± 0.014 | 0.933 | 0.908 | 0.929 ± 0.003 |
| Consensus (majority vote) | 0.704 | 0.688 | 0.720 ± 0.014 | – | – | – |
| *Test set, CD-HIT 0.9* | | | | | | |
| NoiToxML (Full) | 0.654 | 0.636 | 0.650 ± 0.004 | 0.905 | 0.889 | 0.905 ± 0.001 |
| NoiToxML (Boruta) | 0.658 | 0.649 | 0.651 ± 0.004 | 0.903 | 0.896 | 0.903 ± 0.001 |
| NoiToxDP | 0.659 | 0.680 | 0.667 ± 0.007 | 0.920 | 0.911 | 0.920 ± 0.003 |
| NoiToxGP | 0.652 | 0.667 | 0.656 ± 0.013 | 0.910 | 0.907 | 0.908 ± 0.002 |
| Consensus (prob. avg.) | 0.687 | 0.680 | 0.681 ± 0.006 | 0.925 | 0.908 | 0.926 ± 0.001 |
| Consensus (majority vote) | 0.686 | 0.678 | 0.681 ± 0.006 | – | – | – |
| *Independent set, models trained at CD-HIT 0.9* | | | | | | |
| NoiToxML (Full) | 0.694 | 0.651 | 0.694 ± 0.012 | 0.932 | 0.904 | 0.931 ± 0.003 |
| NoiToxML (Boruta) | 0.724 | 0.675 | 0.729 ± 0.008 | 0.919 | 0.904 | 0.921 ± 0.002 |
| NoiToxDP | 0.645 | 0.652 | 0.647 ± 0.020 | 0.900 | 0.903 | 0.901 ± 0.006 |
| NoiToxGP | 0.645 | 0.654 | 0.619 ± 0.022 | 0.891 | 0.883 | 0.882 ± 0.007 |
| Consensus (prob. avg.) | 0.728 | 0.706 | 0.727 ± 0.017 | 0.932 | 0.912 | 0.932 ± 0.002 |
| Consensus (majority vote) | 0.728 | 0.704 | 0.728 ± 0.015 | – | – | – |
